# Compatibility of a competition model for explaining eye fixation durations during free viewing

**DOI:** 10.1101/2025.07.14.664795

**Authors:** Carlos M. Gómez, María A. Altahona, Gabriela Barrera, Elena I. Rodriguez-Martínez

**Affiliations:** Human psychobiology laboratory, Experimental Psychology Department, School of Psychology, University of Seville, Sevilla 41018, Spain

**Keywords:** Eye fixation durations, saccades, competition model, exgaussian model, refractory period

## Abstract

Intersaccadic times or eye fixation durations (EFD) are relatively stable at around 250ms, equivalent to 4 saccades by second. However, the mean and standard deviation are not sufficient to describe the frequency histogram distribution of EFD. The exgaussian has been proposed for fitting the EFD histograms. Present report tries to adjust a competition model (C model) between the saccadic and the fixation network to the EFD histograms. This model is at a rather conceptual level (computational level in Marr’s classification). Both models were adjusted to EFD from an open database with data of 179473 eye fixations. The C model showed to be able, along with exgaussian model, to be compatible for explaining the EFD distributions. The two parameters of the C model can be ascribed to (i) a refractory period for new saccades modeled by a sigmoid equation (A parameter), while (ii) the ps parameter would be related to the continuous competition between the saccadic network related to the saliency map and the eye fixation network, and would be modeled through a geometric probability density function. The model suggests that competition between neural networks would be an organizational property of brain neural networks to facilitate the decision process for action and perception. In the visual scene scanning the C model dynamic justifies the early post-saccadic stability of the foveated image, and the subsequent exploration of a broad space in the observed image. Code to extract the data and to run the model is added at the supplementary material.

## 1 Introduction

The primary function of eye movements is to position objects of interest within the visual field. The reception of visual information is most effective in the fovea, a highly sensitive region of the retina. The ocular motor system is specifically designed to keep objects of interest in focus in this area. Saccadic eye movements permit tracking in a fast manner and focus in the fovea certain parts of the image for intense visual scrutiny during eye fixations.

Saccadic movements are fast and precise, reaching speeds of up to 900°/s in humans. Their purpose is to redirect the gaze efficiently to a new visual stimulus. They have linear relationship between the amplitude and duration of the saccades (Doge and Cline, 1901; Hyde, 1959; Robinson, 1964), as well as between the maximum velocity and the amplitude of the movement (Bahill and Stark, 1979). Due to their high speed, these movements are considered ballistic, i.e., their control depends on a previous calculation of their parameters, without feedback during execution. However, a moderate modulation of trajectory can be obtained in flight by visual inputs (Van Gisbergen and Van Opstal, 1987). Once a saccade is completed, the eyes remain in a fixed position until a new movement is made. During this fixation period, the eyes exhibit small involuntary movements of three different types: tremor, drifting and microsaccades (Carpenter, 1988).

### 1.1 Short description of neural control of saccadic eye movement and eye fixations

The neural control of saccadic movement is complex and implicates a great number of interconnected structures. During saccadic movements directed toward the periphery (abduction), motor neurons and interneurons of the abducens nucleus increase their activity. Conversely, during inward movements (adduction), the firing frequency decreases (Luschei and Fuchs, 1972; Robinson, 1970; Schiller, 1970). This change in activity precedes the movement by approximately 20ms. Once the new ocular position is reached, the firing frequency stabilizes, being higher the more the eye is displaced toward the activation direction. A similar dynamic is recorded in motoneurons of the oculomotor nucleus and the troclear nucleus for movements in the inward/outward and vertical (upper and lower) direction.

The pons reticular formation plays a crucial role in the control of eye movements. Lesions in this region have been observed to cause gaze paralysis (Goebel et al., 1971). Within this area, several neuronal populations have been identified which are related both to the generation of saccadic movements and to the maintenance of the eyes in a fixed position in space. Excitatory and inhibitory burst neurons have a fast and phasic firing. These neurons are organized ipsilateral and contralateral with respect to the oculomotor nuclei to which they project (Luschei and Fuchs, 1972; Van Gisbergen and Robinson, 1977; Van Gisbergen et al., 1981). These neurons are active during eye movements in all directions in space. Excitation and inhibition varies according to the direction of movement. Their main function is to generate the pulse necessary to activate motor neurons and interneurons of oculomotor nuclei, thus allowing the execution of eye movements.

A theoretical model for the generation of saccadic movements has been proposed in which excitatory burst neurons are activated when a saccade is to be made (Van Gisbergen et al., 1981; Robinson, 1981; Zee et al., 1976). These neurons receive information from higher centers about the desired eye position, while they are inhibited by neural information of the eye position calculated by tonic neurons. It has been proposed based on neural network simulations that burst neurons would perform spatial-temporal transformation between the colliculus and the motoneurons, (Smith and Crawford, 2005).

The so-called omnipause neurons exhibit continuous firing in the resting state. However, during a saccade in either direction, they cease their activity until the eye movement concludes (Evinger et al., 1982). These neurons have inhibitory connections with both inhibitory and excitatory burst neurons (Nakao et al., 1980; Evinger and Kaneko, 1977). In addition, their activity can be modulated by visual pathways from the superior colliculus (SC) and optic chiasm, which temporarily inhibits these cells (Kaneko and Fuchs, 1982). Omnipause neurons have the function of inhibiting saccadic movements (Evinger et al., 1977; Evinger et al., 1982). Its dramatic role can be observed by stimulation of omnipause neurons from the rostral tectum which induces its activation, inducing the maintenance of eye position by suppressing saccades, while the activation of the caudal tectum activates the inhibitory burst neurons which liberates the initiation of saccades by inhibiting omnipause neurons (Takahashi and Shinoda, 2018; Takahashi et al., 2022). The saccade stop signal has been attributed not only to omnipause neurons but also to the fastigial nucleus (Rucker et al., 2011).

The nucleus prepositus hypoglossi contains neurons with tonic, tonic-phasic and phasic discharge patterns (Delgado-Garcia et al., 1989). It projects to the ocular motor nuclei and premotor structures such as the cerebellum and vestibular nuclei (Baker et al., 1977; Hikosaka et al., 1977; López-Barneo et al., 1982). The Tonic neurons maintain a continuous and stable firing, the frequency of which is proportional to eye position. These cells play an essential role in maintaining the ocular position once the saccadic movement has been completed. It is proposed that collaterals of excitatory and inhibitory burst neurons connect to the neural integrator which would compute the eye position signal to be maintained by tonic neurons (Robinson, 1981; Delgado-Garcia et al., 1989; Fukushima and Kaneko, 1995). Cerebellar lesion causes eye drift during fixations and is therefore considered to stabilize the “neural integrator” (Robinson, 1975). Not only prepositus hypoglossi presents tonic fixations neurons (Izawa et al., 2004), but also frontal eye fields (FEF), which would contribute to maintain eye fixations by preventing reflexive saccades to other potential targets in the visual field.

The SC stimulation produces rapid eye movements (Adamuck, 1870; Crommelink et al., 1977; Mchaffie and Stein, 1982; McIlwain, 1982; Roucoux et al., 1980), and excitation of oculomotor premotor areas (Grantyn and Grantyn, 1976; Grantyn et al., 1979; Ribas et al., 1983). The SC is considered an important hub for programming saccadic eye movements in which saccade direction is programmed vectorially in a retinotopic map. The strength of functional connectivity from SC to burst neurons is organized in a retinotopic manner (Grantyn et al., 2002; Moschovakis et al., 1998). This pattern of connection permits the transformation from a spatial retinotopic code in the SC into a brainstem temporal code (firing rate and duration) in burst neurons in order to drive extraocular muscles through the motoneurons projection. Saccadic eye movement can be generated without SC by means of connections between FEF and brainstem. However, the SC integrates information from botton-up saliency maps in striate and extrastriate cortex, task requirements from FEF, reward previous history from basal ganglia and substantia nigra, and homeostatic states from zona incerta (reviewed in Veale and Takahashi, 2024 and Takahashi and Veale 2023).

### 1.2 Visual position selection

The top-down and bottom-up influences interacts to generate eye saccadic movements (Rutishauser & Koch, 2007). Bottom-up factors include abruptly occurring stimuli (Yantis & Jonides, 1984), unique features that pop-out in visual search (Treisman & Gelade, 1980), high spatial frequency content and edge density (Mannan, et al.; 1997), higher local spatial contrast (Reinagel and Zador; 1999), luminance contrast and edges (krieger et al., 2000); high spatial frequency edge information (Baddeley & Tatler, 2006); color and others (Frey, König, & Einhäuser, 2007) influence saccadic decisions. These results have permitted to propose the presence of a saliency map, product of the integration of feature maps, which would drive the selection of certain fixation points of the visual scene (Itti & Koch, 2000). Such saliency map would be in interaction with top-down processes for the target selection, with factors such as task demand (Einhäuser, Rutishauser, and Koch ; 2008) and semantic information (Nyström and Holmqvist, 2008).

As outlined above the saccadic system is a very complex system, which on the contrary, when naturally inspecting in a free manner the visual field respond to a relatively simple pattern. We would concentrate here in the temporal aspects, basically in the duration of fixations.

### 1.3 Eye fixation durations (EFD)

During free viewing subjects make around three to four saccades per second, no matter which kind of static image they are looking at (Otero-Millan et al., 2008), or when looking at a video (Chen et al., 2021). The mean of EFD are modulated by experimental conditions as luminance or type of image (Henderson, Nuthmann, & Luke, 2013, Kaspar & König, 2011), but also by the characteristics of next saccade and the experimental task (Unema, et al., 2005, Nuthmann, 2017 ; Schwedes & Wentura, 2016). However, not only first-order statistics as mean and standard deviation are important, but also the probability density function (PDF) that fits the frequency histograms, because the parameters of the distribution would permit to analyze changes in the parameters values related to the experimental conditions as type of images or repetition of images, but also the PDF would suggest some aspects of the internal dynamics that underlies the EFD, interrupted by saccades to new directions. One particular PDF which has been successfully applied to the duration times of fixations is the exgaussian distribution (Guy et al., 2020). As indicated before, to have a PDF with certain parameters, as those of the exgaussian (exponential function convolved with a gaussian distribution to obtain the exgaussian; with parameters µ: mean; σ: standard deviation of the gaussian; and τ: exponent of the exponential function) would permit to identify the possible relationship of these parameters with a defined cognitive processes, or on the contrary its common participation in a given cognitive process (Guy et al., 2020). For instance, for the exgaussian model, more predictable words related to smaller and less lexical ambiguity are related to smaller µ values (Staub, 2011; Sheridan & Reingold, 2012). In the inspection of visual scenes, the exgaussian showed a better fit of EFD histograms than the gaussian distribution (Guy et al., 2020), and the exgaussian parameters were related to specific characteristics of the EFD: the type of image changes the µ Gaussian component, while familiarity was related to the exponential component τ.

### 1.4 Competition model (C model)

The present report tries to frame the EFD in a C model between the saccadic and the fixation system, based on a mutual inhibition and a continuous competition for remaining in the current fixation or producing a saccade to a new location. The model would consider that after a saccade there is a certain refractory period for producing a new saccade. The possibility and generality of such a model has been proved in different settings: eye rivalry, the perception of ambiguous figures (Gómez et al, 1995) and the lever-press response in variable interval schedule of reinforcement (Gómez, 1992). The competition model assumes that the expression of a given percept (or response) occurs when the underlying neural network obtains an activity value higher than that of any other alternative network (Gómez et al., 1992; 1995; 2024), as in the winner-take-all algorithm (Feldman and Ballard; 1982). For eye rivalry and ambiguous percepts would correspond to the winning image representation, and for the lever press would correspond to the competition between the lever press response and any other possible alternative motor response. The model presents the particularity that once a perceptual representation, or just after a response has been made, there is a refractory period for a new perception to be installed or a new response to be produced. This refractory period being modeled by a sigmoid function. Once the asymptotic value of the sigmoid function is reached, a non-biased competition is established (see a more detailed description of the model in the method section). The C model also permits to modulate the probability of a particular percept or response by top-down processes, as for instance attention (Gómez et al., 1995).

Therefore, the successful modeling of the EFD by a competition rule plus a saccadic refractory period would give some hints about the underlying dynamical processes related to maintaining a fixation or the induction of a saccade. It is important to consider here that the exgaussian model has been proved to be a good approximation for fitting the EFD histograms (Guy et al., 2020). And for this reason it would also be fitted to the EFD histograms in present report. However, is not the objective of present report to compare different models, given that different methods for fitting exgaussian and the competition model would be used, but also because other alternative models as the gamma function, the Poisson process and many others would potentially fit the EFD histograms.

A series of complex models which take in account characteristics of the saliency maps (Itti and Koch, 2000), the modelling by random walks for deciding the timing of next saccade (Nuthmann el al., 2010); the neural control of saccadic and eye fixation networks (Findlay and Walker, 1999), or the information gained by stay or go to a new visual scene location (Tatler et al., 2017), have been proposed recently to explain the EFD distributions. The detail of description of these models would be more related to the so-called algorithmic and implementation levels upon Marr’s levels of analysis (Marr, 1982). The simple and parsimonious descriptive C model expressed above, and in the methods section does not try to be compared with such complex models, but to show that from a descriptive manner the EFD distributions can be explained by a PDF with a fixed parameter (A) which define the saccadic refractory period, and a non-stationary time-dependent ps parameter which would be related to the free competition between different positions of the image, once the refractory period for a new saccade has been overcome. In this sense, the model would be more related to the more abstract level of computation in Marr’s proposal (Marr, 1982). The C model possibility to dissociate the often analyzed mean of eye fixation durations in these two parameters (A and ps), defining saccadic refractoriness and steady probability for a new saccade, but also taking in account that it is the competition between neural networks one of the basic process driving EFD, would be the main assets of present model.

Present report tries to show the compatibility of the C model to adjust the EFD histograms in a situation of free viewing of four different type of images presented in five blocks (see methods). These data would be obtained from an open data base (Wilming et al., 2017), which contains the EFD during free viewing of four type of images (nature, urban, fractals and pink noise) during five successive blocks. The proposed hypotheses are that the frequency histograms of EFD are fitted by the C model composed of a (i) geometric PDF modelling the competition between the saccadic and the eye fixation systems, (ii) with the probability of making a saccade in a given time bin (ps) being modulated by a sigmoid to assure the stability of the image for a certain period of time after a saccade.

The model explicitly tests the hypothesis that after a saccade a refractory period for new saccades occurs. Our working hypothesis is that if the sigmoid modulated by the A parameter across very different visual scenes is relatively similar across conditions, it would imply the presence of a post-saccadic refractory motor period. This motor post-saccadic refractory period would permit a deep visual analysis of the foveated region. On the other hand, a variable ps parameter across the different type of presented images, would suggest that the probability of making a saccade would depend on the saliency across the scanned image.

## 2. METHODS

Please notice that in the supplemental materials are the most important scripts (and their functions) used in present report (model simulation, testing the model, creating histograms) and a clear description of how to access the data, and organize them in the MDTM data matrix which is used for analysis. The script order in the supplemental material is suggested to be followed.

### 2.1 Data Base

The data analyzed in this study were obtained from an eye movement and eye fixations data set. This data set stores eye movement recordings and eye fixations from 23 published studies conducted at the Institute of Cognitive Sciences of the University of Osnabruck and the University Medical Center Hamburg-Eppendorf. The data can be obtained from https://datadryad.org/stash/dataset/doi:10.5061/dryad.9pf75, and the general organization of the data are described in Wilmig et al., 2017.

Dataset are stored in an HDF5 file titled “*etdb_v1.0.hdf5*”. This format is extensively used to store large volumes of information in a structured and efficient manner. Each file in the data set contains records organized into vectors that encode information about the fixations. For this analysis, the study entitled ‘Memory I’ was selected. The data and results of this study corresponds to the experiment 1 of Kaspar and König (2011). It contains the following specific information: subject, fixation coordinates on the x and y, fixation start and end expressed in ms, type of figure (nature, urban, fractals, pink noise) number of trial, and block number. The analysis included data from 45 subjects aged between 18 and 48 years, all with normal or corrected-to-normal visual acuity. Before participation, all subjects signed a written consent form to participate in the experiment. During the experiment, participants viewed five blocks of 48 images each with four different categories: nature (12), urban (12), fractals (12) and pink noise (12). Each image was presented for 6 seconds in a random order. The total number of reported fixations was 179473.

As indicated in Kaspar and König (2011), eye tracking was recorded with the Eye Link II system located on a 21” Samsung SyncMaster 1100 CRT monitor. The screen distance was 80 cm and the screen resolution was 1280 × 960 pixels. To facilitate free visual behavior, no headrest was used. Eye movements were recorded at a sampling frequency of 500Hz. Before experimental data recording, subjects were required to perform saccadic movements at different fixation points that appeared on the screen in a randomized order, for eye position calibration purpose.

The detection and characterization of saccadic movements was performed automatically by the eye-tracker, based on three measurements: eye movement of at least 0.1° with a velocity of 30°/s and an acceleration of at least 8000°/s. After the initiation of the saccadic movement, the minimum velocity of saccades was 25°/s and had to be maintained for at least 4ms.

From the data extracted of the data base of Wilmig et al. (2011) we organized the fixation duration as a matrix (matrix name: MDTM) with five columns. Column1: fixation duration. Column 2: Subjects (1-45). Column 3: order of image in each subject (1-240). Column 4: Block number (1-5). Column 5: figure type (nature: labeled as 7; urban:8; fractals: 10, pink noise, 11). A total number of 179473 fixations were analyzed. Each EFD group of fixation inside a block for the same type of images were collapsed and analyzed independently. Therefore, for each subject 20 numerical series of EFD were analyzed in each of the 45 subjects (5 blocks x 4 types of figures). EFD higher than 1.5s were considered outliers and eliminated (total number of outliers= 783, percentage of the total=0.43% of the total number of reported fixations).

### 2.2 Models

The 20 series of EFD (5 blocks x types of images) per subject, were organized in frequency histograms with bins of 50 ms. Then the C model was applied to the EFD histograms. The C model parameters were computed from a homemade function in matlab called from a script.

The C model for the saccadic-eye fixation systems assumes that some sort of competition exists between these two behaviors, without making at this point any hypothesis about the underlying neural mechanism (see the discussion section for that). For the C model the expression of a specific behavior depends on taking a higher neural activity that the alternative behavior (Fig. 1A), following a dynamic similar to the winner-take-all algorithm (Feldman and Ballard, 1982) and the geometric distribution (see Fig. 1B and 1 C, and equation 1). Then, in equation 1 the probability ps would be the probability that the saccade-related network obtains a value greater than the eye fixation controlling network in a time period (bins), and (1-ps) would be the probability that the eye fixation-related network would have an activity value greater than the network controlling saccadic movements in a time bin. For the sake of computation, these probabilities are computed in bins of 50ms. Same duration than the time bins for the built up EFD histograms. Then, the probability (ps) that a particular ocular fixation between two saccades (EFD) get a particular value between time t-1 and time t follows the geometric distribution. The geometric distribution here is applied with the number of time bins (of 50ms) needed for the saccade related network to win the competition with the eye-fixation related network.

**Figure 1.**
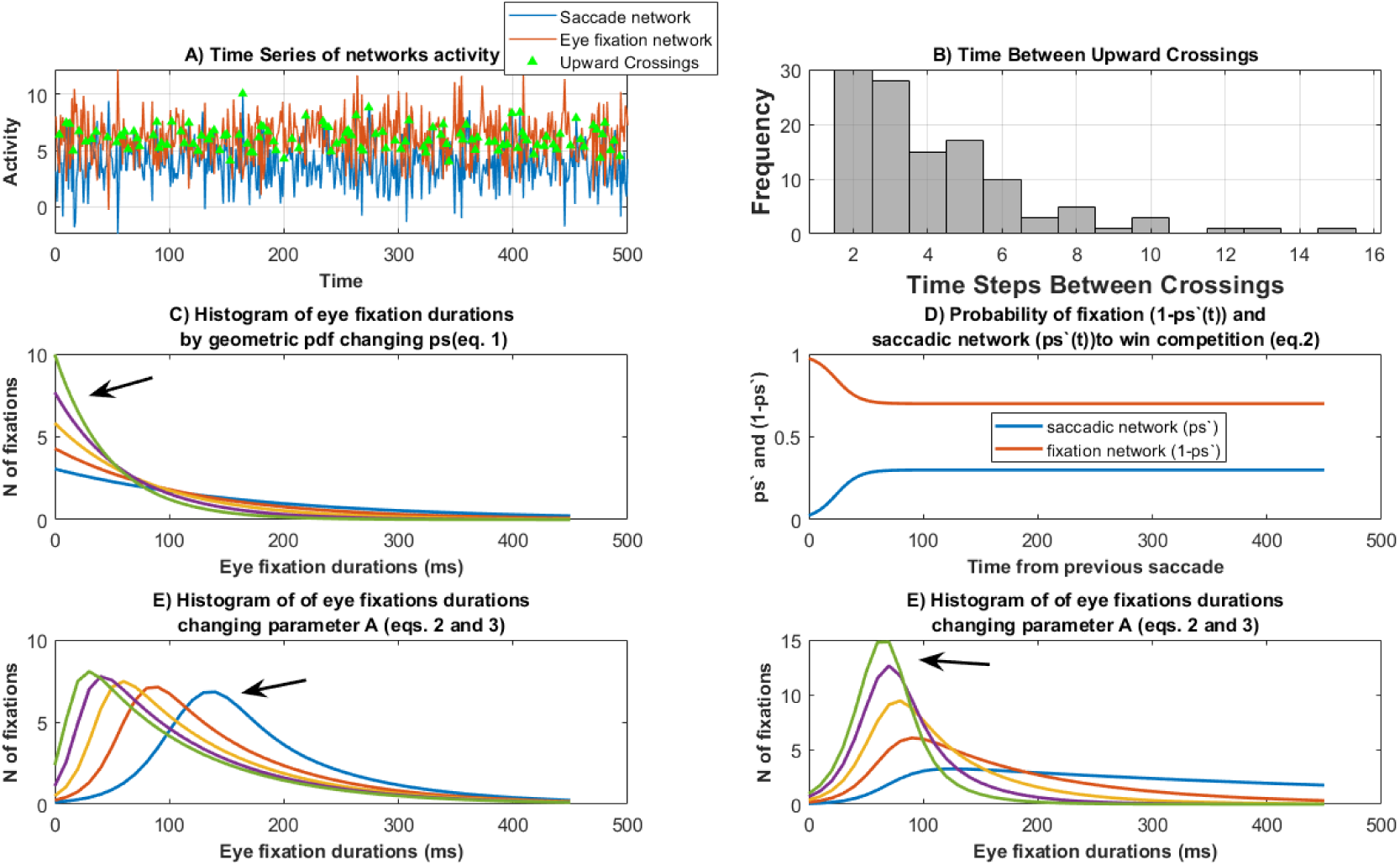
Competition model. A) Simulation of the saccade and eye fixation networks activity. The crossing points in which the saccade network presents more activity than the eye fixation network are labeled. B) Frequency histogram of the time between two crossings of activity higher in the saccade than in the eye fixation network. C) probability density function (PDF) of a geometric distribution following equation 1. The arrow indicates the PDF with the higher ps value. (D) Change in ps’(t) values: probability that the saccade network wins the competition in a given time bin, taking in account the time elapsed from the previous saccade (Equation 2); (1-ps’) represents the probability of the eye fixation network to win the competition. (E) Frequency histogram of fixation duration times as computed from equation 3, obtained by changing the geometric distribution by modulating the A parameter, and consequently the ps’(t) (equation 2). The arrow indicates the PDF with the lower A value. (**F**) Same as (E) but changing asymptotic ps values and keeping fixed the A parameter. The arrow indicates the PDF with the higher ps value.

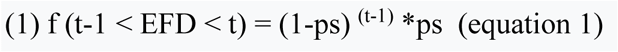

This equation implies that the probability of a fixation time obtaining a certain value (between t-1 and t) depends on the occurrence of the saccade at t=0 and at t, in this sense: f= probability of an intersaccadic value to be between t-1 and t (it can also be expressed as number of fixations by multiplying the PDF by the number of fixations, also in equation 3);

ps = probability that the saccadic network wins;

t= time base (number of time bins from previous saccade needed for the saccadic network to have a higher value than the eye fixation network);

t-1= times the fixation network wins in a row and fixation is maintained

(1-ps)^(t-1)^ = probability that the fixation network wins the competition with the saccadic network in t-1 bins.

Equation 1 implies that the probability of the saccadic network winning the competition (ps) is constant during an eye fixation (or intersaccadic time). The model can be modified to allow ps, initially set at ps=0 for postsaccadic t=0, to increase with time from the last saccade to an asymptotic value of ps. This possibility has been previously successfully assessed in other processes which are also based on a competition rule: The perception duration in eye rivalry and ambiguous figures (Gómez et al., 1995), and the inter-response time of press-lever responses in variable interval schedule of reinforcement (Gómez, 1992). Then, ps would have a relative refractory period in which the probability that a new saccade occurs at a certain time (t), will be a function of time since the last saccade, taking into account that now the probability of the saccadic network is a function of time since the previous saccade (ps’(t)) (Figure 1D and equation 2). For the present C model it is hypothesized that after a given saccade there is an inhibition to a new move after a saccade has been made. For this reason, the probability that the saccadic saliency network wins the competition would a function of the time elapsed from previous saccade (ps’(t)), and would increase as time passes to its asymptotic value (ps):

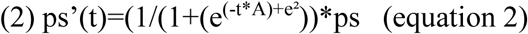

A = parameter to modulate the curvature of the sigmoid.

e² is introduced into the sigmoid equation to have the origin at time zero (ending time of previous saccade).

It must be taken in account that the ps’ parameter dynamics as depicted in Fig. 1D is just the mean value across time points, and it should be considered the mean of an stochastic process (Gómez et al., 2024), which in some time points would permit that ps>(1-ps), and then a saccade would be triggered (Fig. 1A).

Finally, the probability than an EFD gets a given value between t-1 and t under the C model would be (Figures 1E and 1F):

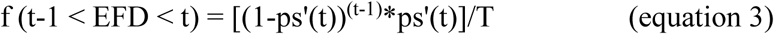

The term T is introduced to normalize equation 3, so that the area under the curve would approximate to 1. T is not computed analytically but numerically, by summing the area under the curve created by the numerator of equation 3.

By changing the curvature parameters of the sigmoid (A), which expresses the dependence of the time elapsed from the previous saccade to the current one, and of the probability that the saccade-related network wins the competition at ps asymptotic levels, different curves can be obtained (See Figure 1E and 1F, respectively). To estimate A and ps, the values of A and ps are changed systematically, and by correlation between the empirical histogram and the equation 3, the optimal A and p parameters are estimated. Once the parameters A and ps have been estimated, the level of significance of the fit between the empirical distribution of EFD and the C model are estimated by means of the Kolmogorov-Smirnoff goodness-of-fit test (Sokal & Rohlf, 1987). The values higher than 700ms were collapsed in EFD histograms and in the model.

As the exgaussian model has proved to be a good approximation for fitting the EFD histograms (Guy et al., 2020), we have reproduced this model and checked its ability to fit the EFD histograms. For that we used the *exgfit* matlab function (Johansson, 2025) to compute the optimal parameters (µ, σ, and τ) of the exgaussian distribution, and then we followed a procedure of optimization of these parameters by changing iteratively the values of these parameters, in a range of +100 to −100 in 1 unit step from the mean values computed with the *exgfit* function (Johansson,2025). The Kolmogorov-Smirnoff goodness-of-fit test (Sokal & Rohlf, 1987) was applied to test the goodness of fit between exgaussian model and the EFD frequency histograms. It is important to consider that as indicated in the introduction section the different fitting procedures used for both distributions (C and exgaussian) don’t permit do claims about superiority of fitting between different models. However, and as an approximation to this issue, the Akaike information criterion (AIC) (Akaike, 1973) was applied to the 900 analyzed EFD histograms (45 subjects x 5 blocks x 4 type of images). The AIC take in account the number of adjusted parameters to decide about the performance of different alternative models. For the application of AIC the exgaussian distribution needed to estimate 3 parameters (mu, sigma and tau), while the competition model needs to adjust the ps value (probability of making a saccade in a defined period of time), and the A parameter which model the refractory period after a saccade has been produced.

### 2.3 Statistical analysis

To test the effects of the factors block and type of image on the mean values of the EFD, and the A and ps parameters of the C model, and ANOVA with these two factors was applied using the JASP 0.19.3.0 (JASP Team, 2024). To test the possible linear relationships between the EFD, the peak time of the of the C model, the peak time of the EFD histogram, and the parameters A and ps of the C model, robust regressions were computed with the *fitlm* function of matlab.

## 3. Results

Figure 2 shows the values of the EFD means for the 5 blocks and 4 type of images. The ANOVA showed that the effect of the block factor (F(2.282,104,42)=4.029; p=0.017;Eta squared=0.014), the type of image factor (F(1.309,57,577)=83.946 ; p<0.001 ; Eta squared=0.458), and the interaction type of image x block were significant (F(6.855,301.606)=2.243 ; p=0.032 ; Eta squared=0.007). The Bonferroni post-hoc for the block factor showed that only the comparison of the block1 with the block 4 presented a trend for significance (p=0.061; Block1<Block4). The post-hoc analysis of the type of image factor showed that the pink noise image presented a higher EFD than all the other type of images (p<0.001, with higher EFD for the pink noise images for the three comparisons). The urban images presented a shorter EFD than the nature and fractal images (p<0.001) (EFD general pattern: urban<nature=fractals<pink noise). There was a complex pattern of interactions between the effects of the block and the images. Given that the purpose of present report is not related to the interpretation of the relationship between blocks and type of image, these interactions are included in table 1 of supplementary materials but they are not further explored.

**Figure 2.**
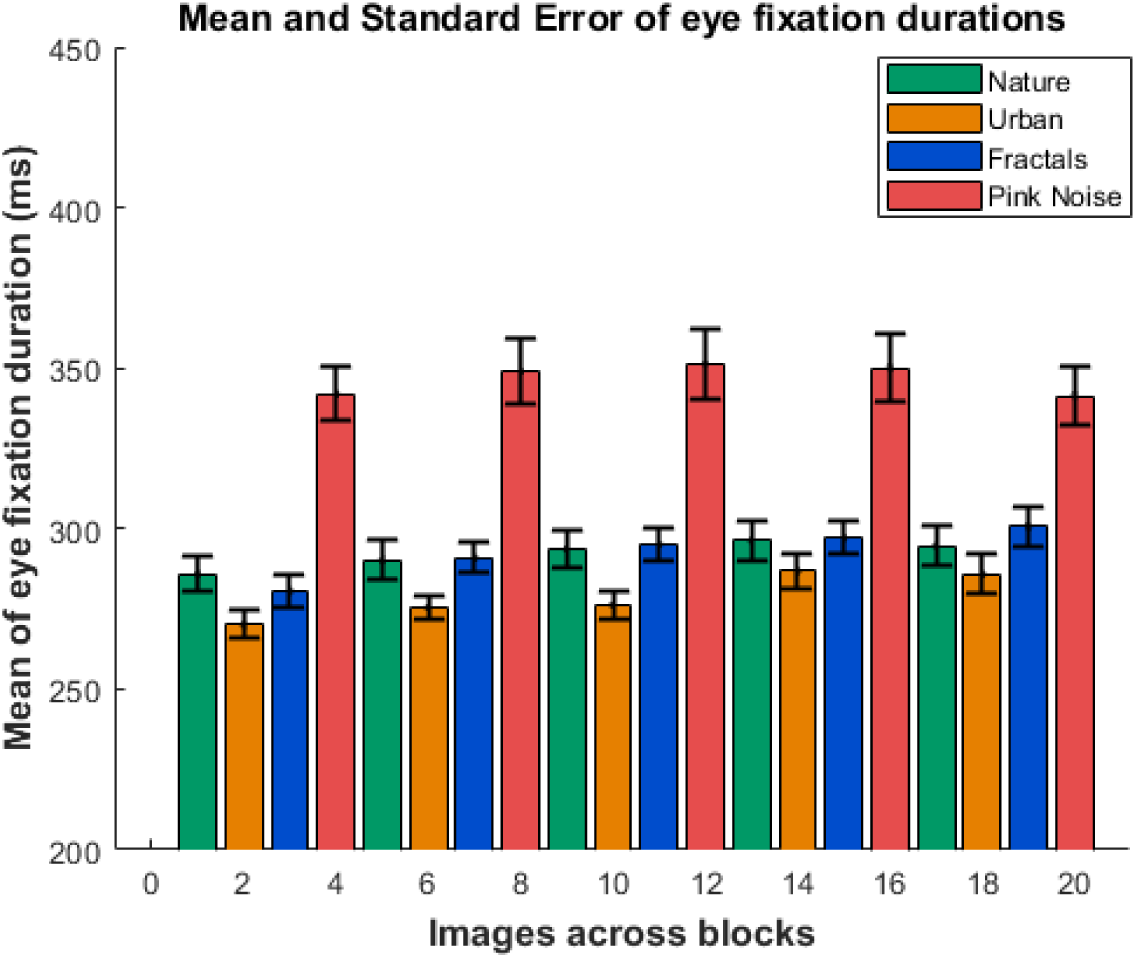
Mean of eye fixation durations. The image shows the mean and standard error of the eye fixation duration across the five consecutive blocks for the four presented type of images (nature, urban, fractal and pink noise).

The Figure 3 shows the overlapping of the C and the exgaussian models on the EFD frequency histograms. Two additional subjects are displayed in the Supplementary figures 1 and 2. The results show that both models are compatible for explaining the EFD histograms. The good fitting of the two models for the EFD histograms can be observed in these figures. The Kolmogorov-Smirnov test of goodness of fit was applied to the 45 subjects, in the 4 type of images and 5 blocks (900 tests), in order to test the adjustment of the competition and exgaussian models to the EFD histograms. Only in 14 cases the exgaussian model did not significantly fitted the EFD histograms. The C model fitted the EFD in all cases. Figure supplementary 3 shows the Akaike information criteria values for the competition and the exgaussian models. The C model showed lower AIC values than the exgaussian model (791 cases over 900 comparisons: 87.9%). However as indicated in the methods section one of the objective of present report is demonstrating the compatibility of both models for explaining the EFD, and given the difficulty of making appropriate comparisons between models due to the different methods used for fitting parameters in both models, no further explorations of this issue were made.

**Figure 3.**
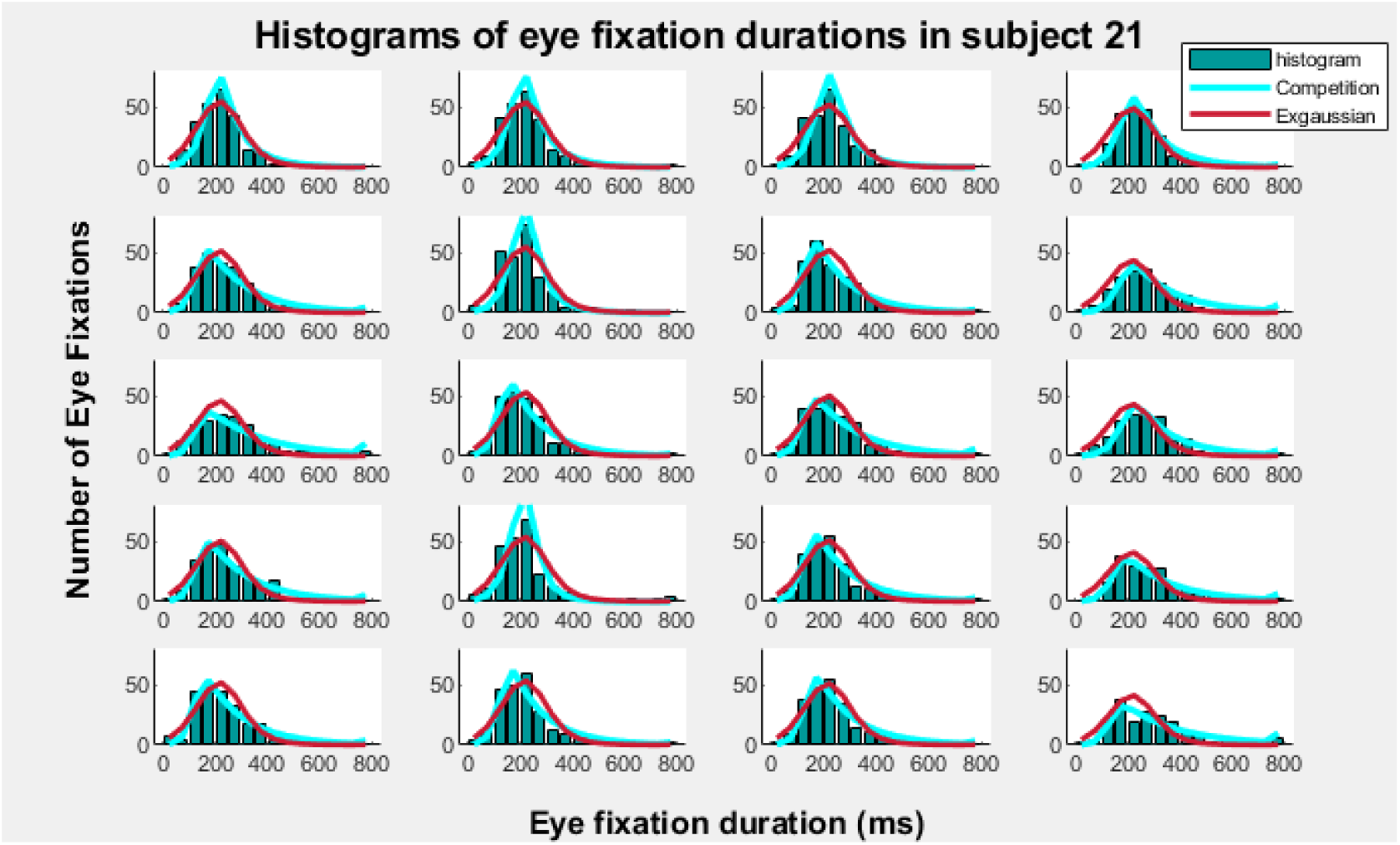
Competition and exgaussian modelling of eye fixation duration histograms. The image shows the fitting of the competition and the exgaussian models of the eye fixation duration frequency histograms for a single subject (subject 21). The fitting of the two distributions is displayed for the four type of images (columns) and for the five block of images presentations (rows).

Fig. 4A shows the mean values for the A parameter of the competition model. The ANOVA showed that only the effect of the block factor was significant (F[3.239,142,505]= 5.30 ; p=0.001 ; Eta squared= 0.031). The bonferroni post-hoc showed differences between block1<block2 (p=0.004), block1<Block3 (p=0.021), and block1<block5 (p=0.034). The results suggest, that given that lower values of A implies a higher refractory period to reach the steady value of the ps parameter, the block1 presented a higher time for reaching the saturation of the C model ps parameter. The absence of an effect of the type of image indicates that the A parameter value was steady across the different images.

**Figure 4.**
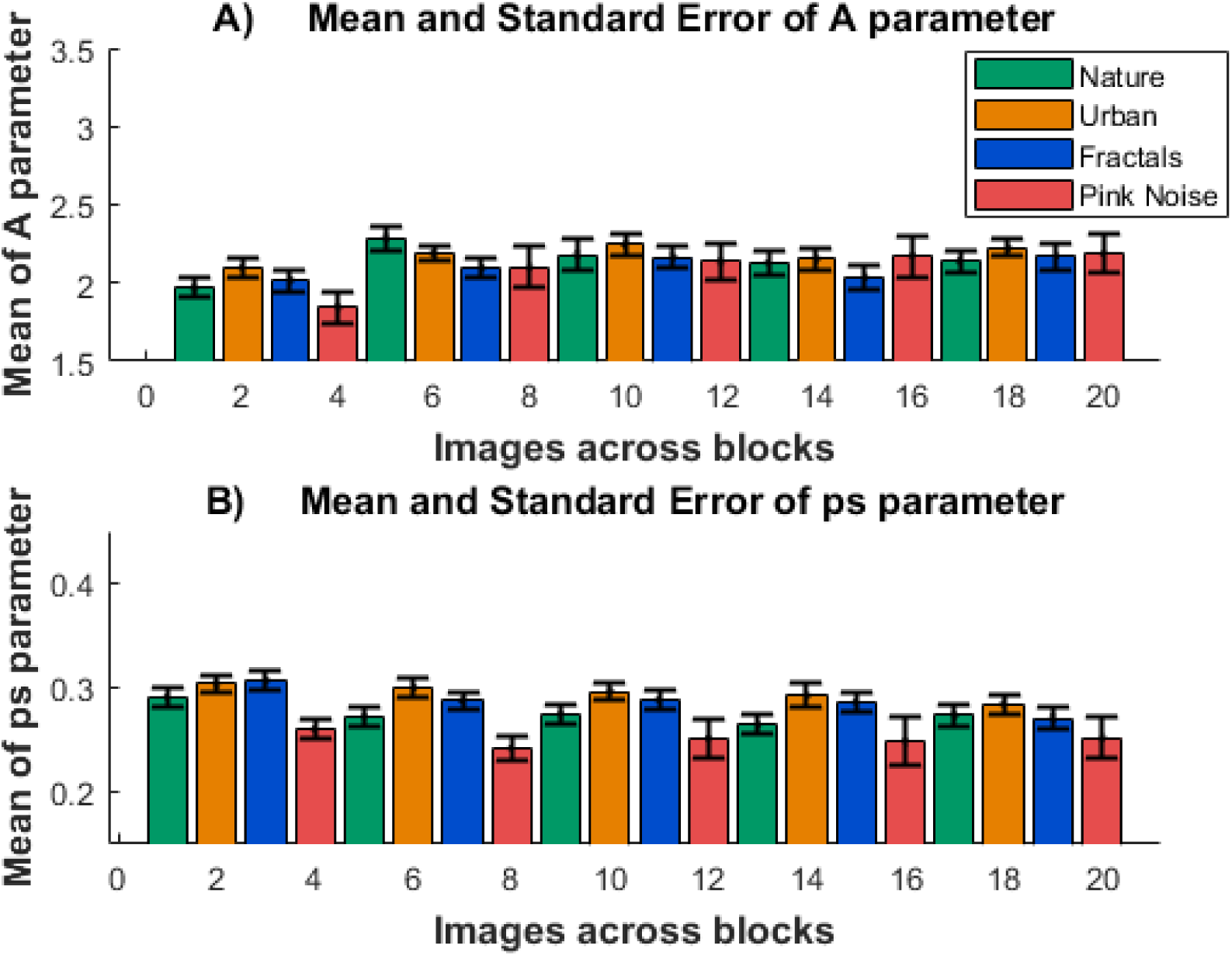
A and ps values of the competition model. A. The image shows the mean and standard error of the A (Fig. 4A) and ps (Fig. 4B) parameters of the competition model, for the five consecutive blocks and for the four presented type of images (nature, urban, fractal and pink noise).

Figure 4B displays the mean values of the ps parameter for the C model. The block factor presented a trend for significance (F[3.019, 132.82 ]= 14.128 ; p=0.073 ; Eta squared=0.011), due to trend for significance of block1>bloc2 (p=0.067). The ANOVA showed that only the effect of the type of image was significant (F[1.879, 82.667 ]= 14.128 ; p<0.001 ; Eta squared=0.065). The bonferroni post-hoc showed differences between the ps parameter for the images: nature<urban (p<0.001); nature<fractals(p=0.047); nature>pink noise(p=0.05); urban>pink noise (p<0.001), fractals >pink noise (p=0.002). The ps parameter results indicate that the probability of doing a saccade in a time period of 50 ms (the period used as bins in the frequency histograms of figure 3), are lower for the pink noise image (ps general pattern: pink noise < nature<urban=fractals).

Figure 5A shows the robust regression between the time to reach the peak time of the C model and the peak of the EFD histograms. The high R^2^ indicate a good adjustment between the model and the empirical data for peak times. Figures 5B and 5C shows the linear robust regression of the A, and ps parameter with the peak time of the EFD histograms, respectively. The much higher R^2^ for the regression of the A parameter suggest that it is the A parameter of the C model which defines the refractory period of the EFD histograms. However, the role of the ps parameter in defining the peak time of the EFD histogram, in a more limited manner, is observed by the statistical significance of the A parameter vs. peak time of the EFD histogram residuals (obtained from 5b), when regressed with the ps parameter (Fig. 5D). The latter results suggest that the empirical peak time of the EFD histogram depends also from the parameter ps, advancing the peak time when ps is high, and inducing a time delay when the ps parameter is low.

**Figure 5.**
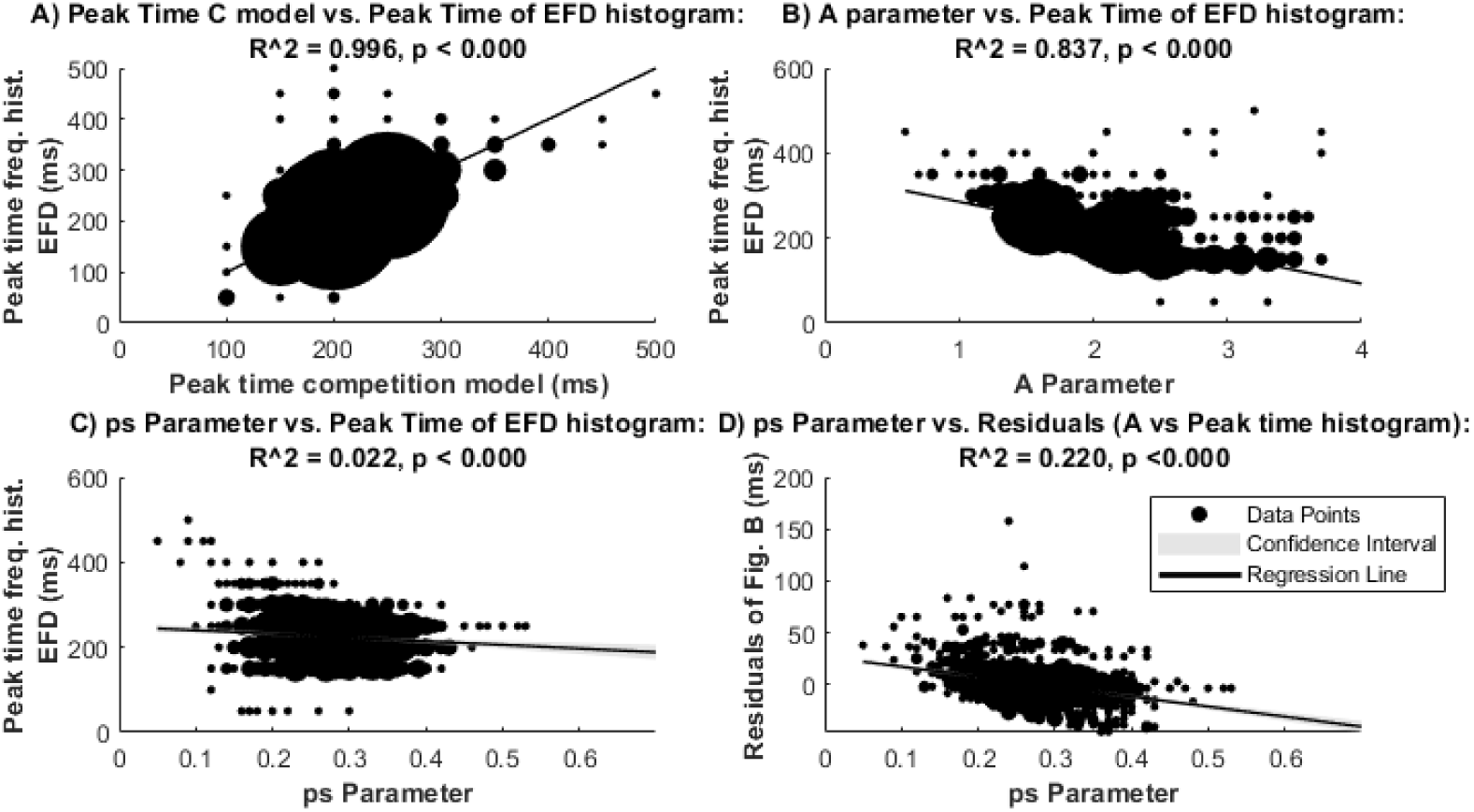
Robust regression between parameters and peak times of the competition model and peak time of the eye fixation duration histograms. Robust regression between the peak time of the empirical peak time of the histograms of the eye fixation durations with the peak time of the competition model (A), the A parameter (B), and the ps parameter (C). 5D shows the residuals of the regression in B vs. the ps parameter. The area of each point is proportional to the number of points with the same value. Confidence intervals of the regression are barely visible due to the high number of represented points (900). EFD: Eye fixation durations.

Figure 6 shows the robust regression of the EFD mean with the A (6A) and ps (6B) of the C model. The higher R^2^ in the ps regression suggests that the ps parameter, indexing the probability of making a saccade in a given interval time, is more related than parameter A for defining the EFD. However, the inverse relationship between the A and ps parameter suggest that both parameters present a certain level of common variance, and therefore for defining the EFD (Fig. 6C).

**Figure 6.**
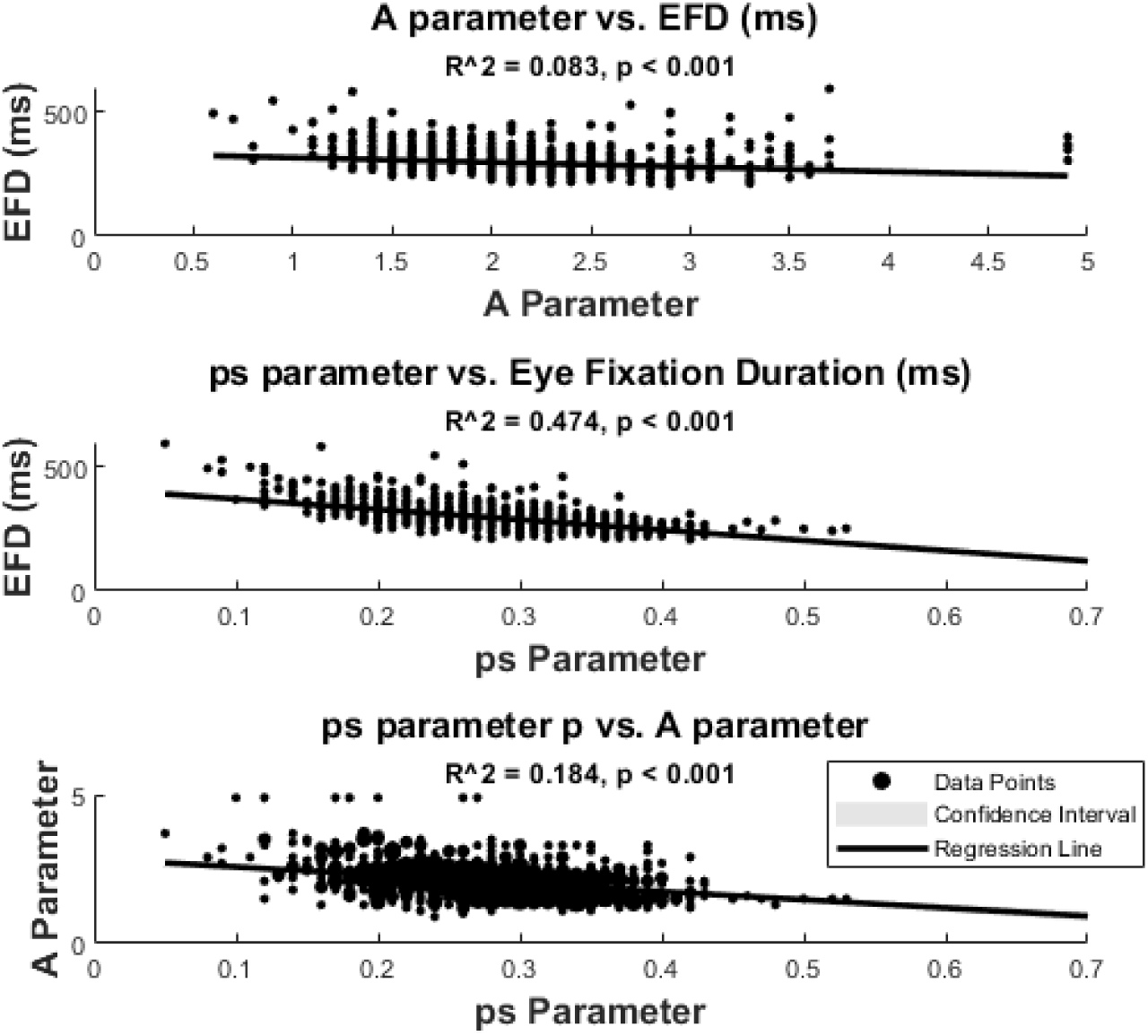
Robust regression of the mean of eye fixation duration (EFD) with A and ps parameters of the competition model. Eye fixation duration regression with the A (6A) and ps (6B) parameters. 6C displays the regression between parameters A and ps. Confidence intervals are barely visible due to the high number of represented points (900).

## 4. Discussion

Present report is based in the same data presented in Kaspar and König (2011), which analyzed the mean of EFD across blocks and type of images. The results obtained in present report broadly replicated the original results, as expected. Kaspar and König (2011) showed an increase in the EFD with the block indicating an increase of EFD for successive presentations. In present report the block factor effect was significant for the EFD (Block1 presenting the shorter EFD across blocks), but bonferroni post-hocs only presented a trend for significance trend in block1 with respect to block 4. For the type of image, the same pattern of urban<nature=fractals<pink noise for the mean of EFD was obtained, same as in Kaspar and König (2011). The authors interpreted these results of increased duration in late presentation blocks as a reflection of a deeper scrutiny of the image in late blocks, while in early blocks the attentional scanning of the images was more related to a more superficial scanning of the image. With respect to the type of image the obtained different durations should be related to the basic characteristics of the image as color or complexity, although possible effects of top-down processes as motivation, aesthetics or other cannot be discarded.

The important point for the present report with respect to the EFD mean analysis, given that we did not try to deepen in the influence of the type of image and block on the EFD, but to test the compatibility of the C model for EFD, is that the general trend obtained for increases of EFD with block and the EFD pattern for the different type of images, is preserved in present analysis when compared with the original analysis (Kaspar and König, 2011). The small differences in the analysis are possibly due to the different methods used to discard outliers. Here, we stablished a very conservative threshold of 1500ms in order to keep a high percentage of eye fixations to give robustness to the model fitting. The later procedure produced a higher number of accepted fixations than in Kaspar and König (2011), which used a limit of two standard deviations as a limit. The threshold limit in present report also produces higher EFD means than in the original work.

The importance of defining not only first-order statistics of the EFD data has been highlighted by Guy et al. (2020). They have indicated that the use of central or dispersion measures can only account for a general picture of the data distribution, given that similar values can be obtained with a very different structure of the data. The exgaussian model, in which a gaussian distribution is convolved with an exponential distribution has been broadly applied to explain the EFD histograms. This model has been proved to be a good approximation to explain the EFD histograms (Guy et al., 2019; Glaholt, Rayner, & Reingold, 2013; Luke et al., 2014). The results of present report also showed that the exgaussian model is compatible with the empirical EFD data from the Kaspar and König report (2011). Furthermore, the exgaussian model has been successfully applied to explain EFD histograms in reading tasks (Staub, 2011; Sheridan & Reingold, 2012) ; in the inspection of visual scenes (Guy et al., 2020), permitting to characterize the exgaussian parameters to the type of image and familiarity. Kieffaber et al. (2006) proposed the parameters defining the exgaussian would be related to different psychological processes, as µ to perceptual processes and τ to decision processes. The importance of the exgaussian parameters as metrics to be correlated with cognitive processes is without no doubt an interesting approach, which would permit to associate the parameters to specific computational brain processes.

On the other hand, the structure of the C model (Gómez, 1992; Gómez et al., 1995; Gómez et al., 2024) makes two very specific proposals with respect to the parameters characterizing it, (i) there is an asymptotic probability to the value of making a saccade in a given time bin (ps), but (ii) after a saccade there is a reduced probability of making a saccade, which is modeled by a sigmoid dependent of the A parameter. The C model was validated by the goodness of fit with the EFD histograms, but also by the very high correlation between the peak times of the model and the peak time of the EFD histograms. In present report the Akaike information criteria (Akaike,1973) assigns a better performance to explain the EFD to the C with respect to the exgaussian model. However, the different methods used to adjust the parameters of both models make it difficult to make an appropriate comparison between the two models. The main point for these comparisons in present report is that given the very high frequency of cases in which both models fitted the EFD histograms, both are numerically compatible for explaining the EFD. However, we would discuss only the possible meaning of the C model, main objective of present report.

The A parameter of the C model, showed a high correlation with the peak time of the EFD histogram and with the peak time of the histograms of the C model, indicating that A it is responsible for modelling the refractory time period after a saccade. The A parameter showed a lower value in the block1 which implies a longer time to reach the peak time in block 1. The main role of the time to reach the histogram peak rather than to the total duration of the EFD, is supported by the modest relationship between the A parameter and EFD. The ANOVA showed that the A parameter is relatively steady across blocks and type of images (see below for A as a possible index of a fixed motor refractory period). On the other hand, the ps parameter in the C model defines the probability of making a saccade in a time bin (50 ms in present report), once the transient refractory period for making a saccade is over. Its close relationship with EFD appears in the similar pattern (although inverted) of post-hoc comparisons between EFD means and ps means for the type of images: EFD general pattern is urban<nature=fractals<pink noise; and ps general pattern: pink noise < nature<urban=fractals). Furthermore, the ps high relationship with EFD, much higher that the relationship between the A parameter and the EFD, suggests its critical role for defining the EFD. A closer look shows that the residuals of the relationship between histogram peak time vs. A, are related to ps, suggesting that the peak time of the histogram depends on both: primarily on the A parameter defining the refractory period, but modulated by ps defining the probability of making a saccade, or (1-ps) staying in the current position (see Figure 1F). The latter is important, because we would discuss later that the preferential time for looking at the current visual scene position (time until the peak of the EFD histogram), is related primarily to a relatively fixed A parameter, but also to a more variable ps parameter which would modulate the time to peak of EFD histograms.

As any model in which external data are interpreted from an internal model, which is not empirically and simultaneously measured, it can be defined as a modeling inverse problem. The latter argument implies that the empirical EFD could be explained by different internal models. Additional information about the internal model would help in increasing its validity. Therefore, some comments should be done about the possibility that the A and ps parameter would be related to some neuroanatomical structures and neural dynamics of the oculomotor and saliency networks, that although not here empirically recorded would suggest a neural compatibility with the C model for explaining EFD.

The SC plays an essential role in defining the amplitude, direction but also for deciding when to move the eyes, equivalent to ending eye fixation, although other areas as the FEF can also initiate saccades (Grantyn et al., 2002; Moschovakis et al., 1998; Veale and Takahashi, 2024;Takahashi and Veale 2023). The SC activates the brainstem oculomotor plant by increasing the activity of burst neurons (Van Gisbergen et al., 1981), and inhibiting the omnipause neurons (Evinger et al., 1982; Takahashi et al., 2022). The role of the SC for selecting the direction of movements under competitive conditions has been demonstrated in a two choice experimental situation, in which excitatory bias from FEF would permit a higher activity in the positively biased colliculus, which would be selected for action through a winner-take-all process, producing contraversive eye movements (Lintz et al., 2019). Such model could be generalized to human visual scanning and EFD distributions as those presented in this report. One critical aspect of the here presented C model is the need for inhibition in the area generating the saccade, something that occurs internally in the SC, but also from the projection to colliculus from substantia nigra pars reticulata (Hikosaka and Wurtz,1989). SC should receive information from the saliency maps (Itty and Koch, 2000). The saliency map implies also some sort of center-surround inhibition in the different representational maps created from bottom-up information and modulated by top-down inputs representing task, motivation and many other possible aspects. The different saliency maps representing visual features as luminance contrast, color, opponency, oriented edges, flicker and motion, should be finally integrated for generating a priority signal that would control orienting behavior, and optionally in overt attention a saccadic movement. Saliency maps and priority maps have been described in several brain areas as V1, V4; lateral intraparietal area, FEF, dorsolateral prefrontal cortex and SC (reviewed in White et al., 2017). If neuroanatomy, including the presence of inhibitory synaptic connection, can be considered compatible, the dynamic aspects are much more difficult to be interpreted from the current scientific literature. It can be suggested that the refractory period for making a new saccade after completion of the previous saccade should be due to an inhibition of the motor programs to produce a saccade, and on the other hand the possibility that inhibition of the of the current fixated position in saliency map of the visual scene occurs, which is being progressively adapted permitting other areas of the saliency maps to win competition for gaze orienting in a winner-take-all type competition (Feldman et al., 1982; Gómez, 1992; Gómez et al., 1995).

From the modelling to neural implementation two different subprocesses can be suggested: A motor refractory period indexed by the A parameter, relatively steady, and a free competition period to generate a saccade which would be indexed by a more variable ps parameter very dependent of the image type. However, the peak time of EFD histograms would also be influenced by the ps parameter, with higher ps values shortening the peak time and low values increasing the peak time. The presence of a refractory period in the immediate post-saccadic period has been previously reported, and interpreted as result of a random exponential model triggering saccades in the so called α phase of the model (Harris et al., 1988). The refractory period has also been modeled by a Poisson process constrained by a Gaussian inhibition period (refractory period), and a follow-up rebound of the Poisson λ parameter (Amit et al., 2017). Both model shares with the C model the proposal of a relatively fixed refractory period duration, that would influence the relatively steady rate of saccades, as it has also proposed for other perceptual systems with fixed sensory inputs (Gómez et al., 1995). One possible source of this refractory period would be related to the so-called saccadic inhibition phenomenon, in which a saccade is delayed if a peripheral target is presented. Saccadic inhibition, has been proposed to relay in the post-saccadic activity in visual cortex, then rooted to transiently inhibit the oculomotor system (Amit et al., 2017). The main difference with the C model is for the phase of free competition, given that the two indicated models relies in random exponential or Poisson processes (Harris et al., 1988; and Amit et al., 2017), while the C model relies in the neurophysiological plausible concept of push-pull competition with a dynamics similar to a winner-take-all mechanism (levels 3 and 2 of Findlay and Walker, 1999). Although present model is still stochastic, as the ps parameter is the mean probability of making a saccade in a given time. and (1-ps) would be the mean probability to maintain fixation in the same period, it can be easily rooted in the widespread activation-inhibition processes of the brain. If the decision to make a saccade is finally mainly taking place in the retinotopic map, the where and when to look would rely in competition not only in the colliculus but in different cortical areas related to oculomotor and saliency maps (Veale and Takahashi, 2024; and Takahashi and Veale, 2023). This competition would depend of possible top-down influences, of the familiarity acquired during the experiment, and from the characteristics of the images (Kaspar and König 2011), something which is would be captured by the parameter ps, whose value would depend on the features described in the introduction section (Yantis & Jonides, 1984; Treisman & Gelade, 1980, Mannan, et al., 1997; Reinagel and Zador, 1999; Krieger et al., 2000; Baddeley & Tatler, 2006; Frey, König, & Einhäuser, 2007).

A final point to be discussed is if the empirical EFD transient refractory period could be considered a fixed period, possibly related to post-saccadic motor inhibition with a very rigid time dynamics, or on the contrary, would be a consequence to a progressive adaptation to the current scrutinized visual scene position, which would be more variable and dependent of image characteristics. The post-saccadic motor inhibition hypothesis is suggested by the relatively low differences in values of the A parameter across type of images, although the much more variable ps parameter is also influencing the peak time of the EFD histograms obtained empirically, justifying the variability in peak time of the empirical EFD histogram and of the peak time of the model fitting. Those results suggest that the refractory period of empirically recorded EFD would occur as a consequence of both: a relatively fixed post-saccadic motor inhibition similar to that described for the saccadic inhibition phenomenon (Amit et al, 2017), plus an influence of the image features, which would delay or advance the empirical refractory period. The significant differences of the ps parameter across type of images, would suggest that the saliency of the image would influence the probability of inducing a saccade. Once, the sigmoid defining the ps parameter saturates, the free competition at the saliency map would be allowed and would be modeled by a geometric distribution. The competition side of the C model, given the stochastic character of ps’, would permit the exploration of a broad content in the observed image, and don’t get stuck in a given position of the image.

## Supporting information

figures, statistics, matlab code

## Data and code availability

The data analyzed in this study were obtained from an eye movement data set. This data set stores eye movement recordings from 23 published studies conducted at the Institute of Cognitive Sciences of the University of Osnabruck and the University Medical Center Hamburg-Eppendorf. The data can be obtained from https://datadryad.org/stash/dataset/doi:10.5061/dryad.9pf75, and the general organization of the data are described in Wilmig et al., 2017. The code can be provided upon a reasonable request

## Funding

This research was funded by Agencia Estatal de Investigación, grant number PID2022-139151OB-I00

## Authors and Affiliations

Carlos M. Gómez, María A. Altahona. Grabriel barrera and Elena I. Rodriguez-Martinez Lab of Psychobiology, Dept of Experimental Psychology, University of Sevilla, c/Camilo José Cela s/n, Sevilla, 41018 Spain

## Contributions

Carlos M. Gómez: Formal Analysis, Investigation, Methodology, Software, Validation writing; María A. Altahona: Methodology, Visualization, Writing; Gabriela Barrera: Visualization, writing, scientific search and review; Elena I. Rodríguez-Martínez: Visualization, writing, scientific search and review, supervision.

## Ethics declarations

### Conflict of interest

The authors declare that they have no conflict of interest.

### Ethics approval and consent to participate

No ethical approval needed. Data were from an open database.

### Consent for publication

No extra consent for publication needed.

## Notes

### Competing Interest Statement

The authors have declared no competing interest.

### Summary of Updates

In the abstract we have included the text: Code to extract the data and to run the model is added at the supplementary material. In the methods: Please notice that in the supplemental materials are the most important scripts (and their functions) used in present report (model simulation, testing the model, creating histograms) and a clear description of how to access the data, and organize them in the MDTM data matrix which is used for analysis. The script order in the supplemental material is suggested to be followed. In the results we have changed: the number 8 by the number 14 (feferingg to the nimber of cases the exgaussian model did not fit the data). In the supplemental material we have added the statistical contrasts and the matlab code

## REFERENCES

1. Adamück, E (1870). Über die Innervation der Augenbewegungen. Zentralbl. Med. Wiss. 8, 65–67.

2. Akaike, H (1973). Information theory and an extension of the maximum likelihood principle. In Petrov BN. & Csaki BF. (Eds.), Second International Symposium on Information Theory (pp. 267–281). Academiai Kiado: Budapest.

3. Amit R, Abeles D, Bar-Gad I, Yuval-Greenberg S (2017).Temporal dynamics of saccades explained by a self-paced process. Sci Rep. 7(1):886 doi:10.1038/s41598-017-00881-7

4. Baddeley RJ, Tatler BW (2006). High frequency edges (but not contrast) predict where we fixate: A Bayesian system identification analysis. Vision Res. 46(18):2824–2833. doi:10.1016/j.visres.2006.02.024

5. Baker R, Berthoz A, Delgado-García J (1977). Monosynaptic excitation of trochlear motoneurons following electrical stimulation of the prepositus hypoglossi nucleus. Brain Res. 121:157–161. doi:10.1016/0006-8993(77)90445-0

6. Bahill AT, Stark L(1979). The Trajectories of Saccadic Eye Movements. Scientific American. 240:108–117.

7. Carpenter RHS (1988). Movements of the eyes. Pion, London.

8. Chen CY, Matrov D, Veale R, et al. (2021). Properties of visually guided saccadic behavior and bottom-up attention in marmoset, macaque, and human. J Neurophysiol. 125(2):437–457. doi:10.1152/jn.00312.2020

9. Crommelink M, Guitton D, Roucoux A (1977). Retinotopic versus spatial coding saccades: clues obtained by stimulating a deep layer of cat’s superior colliculus. In: Baker R, Berthoz A (eds) Control of Gaze by Brain Stem Neurons. Elsevier North-Holland, pp. 425–435.

10. Delgado-García JM, Vidal PP, Gómez C, Berthoz A (1989) A neurophysiological study of prepositus hypoglossi neurons projecting to oculomotor and preoculomotor nuclei in the alert cat. Neuroscience. 29:291–307. doi:10.1016/0306-4522(89)90058-4

11. Dodge R, & Cline TS (1901). The angle velocity of eye movements. Psychol. Rev., 8(2), 145–157. 10.1037/h0076100

12. Einhäuser W, Rutishauser U, Koch C (2008). Task-demands can immediately reverse the effects of sensory-driven saliency in complex visual stimuli. J Vis. 8(2):2.1–19. doi: 10.1167/8.2.2.

13. Evinger C, Kaneko C.R.S, Johansen G. W and Fuchs AF, Omnipause cells in the cat (1977). In Baker y A. Berthoz (Eds.) Control of Gaze by Brain Stem Neurons, Elsevier / North Holland Biomedical Press. pp 337–348

14. Evinger C, Kaneko CR, Fuchs AF (1982). Activity of omnipause neurons in alert cats during saccadic eye movements and visual stimuli. J Neurophysiol. 827–844. doi:10.1152/jn.1982.47.5.827

15. Feldman J, & Ballard D, (1982). Connectionist models and their properties. Cogn Sci. 6, 205–254.

16. Findlay JM, Walker R. A (1999).model of saccade generation based on parallel processing and competitive inhibition. Behav Brain Sci. 22(4):661–721. doi:10.1017/s0140525x99002150

17. Frey HP, König P, Einhäuser W (2007).The role of first- and second-order stimulus features for human overt attention. Percept Psychophys. 69(2):153–161. doi:10.3758/bf03193738

18. Fukushima K, Kaneko CR. Vestibular integrators in the oculomotor system. (1995). Neurosci Res. 22(3):249–58. doi: 10.1016/0168-0102(95)00904-8.

19. Glaholt MG, Rayner K, Reingold EM (2013). Spatial frequency filtering and the direct control of fixation durations during scene viewing. Atten Percept Psychophys. 75(8):1761–1773. doi:10.3758/s13414-013-0522-1

20. Gómez C, Argandoña ED, Solier RG, Angulo JC, Vázquez M. Timing and competition in networks representing ambiguous figures. Brain Cogn. (1995) 29(2):103–114. doi:10.1006/brcg.1995.1270

21. Gómez CM, Rodríguez-Martínez EI, & Altahona-Medina MA, (2024) Unavoidability and Functionality of Nervous System and Behavioral Randomness. Appl Sci. 14(10), 4056. 10.3390/app14104056

22. Gómez CA, (1992).Competition Model of IRT Distributions During the First Training Stages of Variable-Interval Schedule. Psychol Rec 42, 285–293 10.1007/BF03399602

23. Goebel HH, Komatsuzaki A, Bender MB, Cohen B (1971). Lesions of the pontine tegmentum and conjugate gaze paralysis. Arch Neurol. 24:431–440. doi:10.1001/archneur.1971.00480350065007

24. Grantyn AA, Grantyn R (1976). Synaptic actions of tectofugal pathways on abducens motoneurons in the cat. Brain Res. 105(2):269–285. doi:10.1016/0006-8993(76)90425-x

25. Grantyn A, Grantyn R, Robiné KP, Berthoz A (1979). Electroanatomy of tectal efferent connections related to eye movements in the horizontal plane. Exp Brain Res. 37(1):149–172. doi:10.1007/BF01474261

26. Grantyn A, Brandi AM, Dubayle D, Graf W, Ugolini G, Hadjidimitrakis K, Moschovakis A (2002). Density gradients of trans-synaptically labeled collicular neurons after injections of rabies virus in the lateral rectus muscle of the rhesus monkey. J Comp Neurol. 451(4):346–61.

27. Guy N, Azulay H, Kardosh R, et al. (2019). A novel perceptual trait: gaze predilection for faces during visual exploration. Sci Rep. 9(1):10714. doi:10.1038/s41598-019-47110-x

28. Guy N, Lancry-Dayan OC, Pertzov Y (2020). Not all fixations are created equal: The benefits of using ex-Gaussian modeling of fixation durations. J Vis. 20(10):9. doi: 10.1167/jov.20.10.9.

29. Harris CM, Hainline L, Abramov I, Lemerise E, Camenzuli C (1988). The distribution of fixation durations in infants and naive adults. Vision Res.28(3):419–432. doi:10.1016/0042-6989(88)90184-8

30. Hikosaka O, Kawakami T (1977). Inhibitory reticular neurons related to the quick phase of vestibular nystagmus--their location and projection. Exp Brain Res. 27:377–386. doi:10.1007/BF00235511

31. Henderson JM, Nuthmann A, Luke SG (2013). Eye movement control during scene viewing: immediate effects of scene luminance on fixation durations. J Exp Psychol Hum Percept Perform 39(2):318–322. doi:10.1037/a0031224

32. Hikosaka O, Wurtz RH.(1989) The basal ganglia. Rev Oculomot Res. 3:257–81. PMID: 2486325

33. Hyde J.E. (1959). Some Characteristics of voluntary human ocular eye movements in the horizontal plane. Am. J. Ophtalmol. 48: 85–94.

34. Itti L, Koch C (2000). A saliency-based search mechanism for overt and covert shifts of visual attention. Vision Res. 40(10-12):1489–1506. doi:10.1016/s0042-6989(99)00163-7

35. Izawa Y, Suzuki H, Shinoda Y (2004). Suppression of visually and memory-guided saccades induced by electrical stimulation of the monkey frontal eye field. I. Suppression of ipsilateral saccades. J Neurophysiol. 92(4):2248–60. doi: 10.1152/jn.01021.2003. PMID: 15381744.

36. JASP Team(2024). JASP (Version 0.19.3) [Computer software).

37. Johansson, T (2025). exgfit - Fit ExGaussian distribution to data (https://www.mathworks.com/matlabcentral/fileexchange/70225-exgfit-fit-exgaussian-distribution-to-data), MATLAB Central File Exchange.

38. Kaneko CR, Fuchs AF (1982). Connections of cat omnipause neurons. Brain Res. 241:166–170. doi:10.1016/0006-8993(82)91240-9

39. Kaspar K, König P (2011).Overt attention and context factors: the impact of repeated presentations, image type, and individual motivation. PLoS One. 6(7):e21719. doi:10.1371/journal.pone.0021719

40. Kieffaber PD, Kappenman ES, Bodkins M, Shekhar A, O’Donnell BF, Hetrick WP (2006). Switch and maintenance of task set in schizophrenia. Schizophr Res. 84(2-3):345–358. doi:10.1016/j.schres.2006.01.022

41. Krieger G, Rentschler I, Hauske G, Schill K, Zetzsche C. (2000) Object and scene analysis by saccadic eye-movements: an investigation with higher-order statistics. Spat Vis. 13(2-3):201–214. doi:10.1163/156856800741216

42. Lintz MJ, Essig J, Zylberberg J, Felsen G. (2019) Spatial representations in the superior colliculus are modulated by competition among targets. Neuroscience. 408:191–203. doi: 10.1016/j.neuroscience.2019.04.002

43. Mannan SK, Ruddock KH, Wooding DS (1997). Fixation patterns made during brief examination of two-dimensional images. Perception. 26(8):1059–1072. doi:10.1068/p261059

44. Marr, D. (1982). Vision: A Computational Approach, San Francisco, Freeman & Co.

45. Mchaffie JG, and Stein BE, (1982). Eye movement evoked by electrical stimulation in the superior colliculus of rats and hamsters. Brain Res. 247: 243–253.

46. McIlwain JT (1982). Lateral spread of neural excitation during microstimulation in intermediate gray lager of cat’s superior colliculus. J. Neurophysiol. 47: 167–178

47. Moschovakis AK, Kitama T, Dalezios Y, Petit J, Brandi AM, Grantyn AA (1998). An anatomical substrate for the spatiotemporal transformation. J Neurosci. 18(23):10219–10229. doi:10.1523/JNEUROSCI.18-23-10219.

48. Nakao S, Curthoys IS, Markham CH (1980). Direct inhibitory projection of pause neurons to nystagmus-related pontomedullary reticular burst neurons in the cat. Exp Brain Res. 283–293. doi:10.1007/BF00237793

49. Nuthmann A, Smith TJ, Engbert R, Henderson JM (2010). CRISP: a computational model of fixation durations in scene viewing. Psychol Rev. 117(2):382–405. doi:10.1037/a0018924

50. Nuthmann A (2017). Fixation durations in scene viewing: Modeling the effects of local image features, oculomotor parameters, and task. Psych Bull & Rev. 24(2), 370–392.

51. Nyström M, Holmqvist K(2008). Semantic Override of Low-Level Features in Image Viewing–Both Initially and Overall. J. Eye Mov. Res. 2(2):1–11. 10.16910/jemr.2.2.2

52. Lopez-Barneo J, Darlot C, Berthoz A, Baker R (1982). Neuronal activity in prepositus nucleus correlated with eye movement in the alert cat. J Neurophysiol. 47(2):329–52. doi: 10.1152/jn.1982.47.2.329.

53. Luschei ES, Fuchs AF (1972). Activity of brain stem neurons during eye movements of alert monkeys. J Neurophysiol. 35(4):445–461. doi:10.1152/jn.1972.35.4.445

54. Luke SG, Smith TJ, Schmidt J, Henderson JM (2014). Dissociating temporal inhibition of return and saccadic momentum across multiple eye-movement tasks. J Vis. 14(14):9

55. Otero-Millan J, Troncoso XG, Macknik SL, Serrano-Pedraza I, Martinez-Conde S (2008). Saccades and microsaccades during visual fixation, exploration, and search: foundations for a common saccadic generator. J Vis. 8(14):1–18.

56. Ribas J, Serra R, Álvarez de Toledo G (1983). Potenciales postsinápticos en estimulación del Mns del núcleo del III colículo superior. In: Actas de la Reunión de la III Neurobiólogos Españoles, pp 114.

57. Robinson DA (1964.) The Mechanics of Human Saccadic Eye Movements. J Physiol. 174: 245 –264.

58. Robinson DA (1970). Oculomotor behavior in the monkey. J Neurophysiol. 33: 393–404.

59. Robinson DA (1975). Oculomotor control signals. In Lennestrand (Eds.) Basic Mechanisms of ocular Motility and their Clinical Implications, Pergamon, Turkey P. Bach-y-Rita u. G., pp 337–374

60. Robinson DA (1981). The use of control systems analysis in the neurophysiology of eye movements. Annu Rev Neurosci. 463–503. doi:10.1146/annurev.ne.04.030181.002335

61. Reinagel P, Zador AM. (1999). Natural scene statistics at the center of gaze. Network 10(4):341–350.

62. Roucoux A, Guitton D, Crommelinck M (1980). Stimulation of the superior colliculus in the alert cat. Exp Brain Res. 39:78–85.

63. Rutishauser U, Koch C (2007). Probabilistic modeling of eye movement data during conjunction search via feature-based attention. J Vis.7(6):5. doi:10.1167/7.6.5

64. Rucker JC, Ying SH, Moore W, Optican LM, Büttner-Ennever J, Keller EL, Shapiro BE, Leigh RJ (2011). Do brainstem omnipause neurons terminate saccades? Ann N Y Acad Sci. 1233:48–57. doi: 10.1111/j.1749-6632.2011.06170.x.

65. Schiller PH (1970). The discharge characteristics of single units in the oculomotor and abducens nuclei of the unanesthetized monkey. Exp Brain Res. 10:347–362. doi:10.1007/BF02324764

66. Schwedes C, Wentura D (2016). Through the eyes to memory: Fixation durations as an early indirect index of concealed knowledge. Mem Cognit. 44(8):1244–1258. doi: 10.3758/s13421-016-0630-y.

67. Sheridan H, Reingold EM (2012). The time course of contextual influences during lexical ambiguity resolution: evidence from distributional analyses of fixation durations. Mem Cognit. 40(7):1122–31. doi: 10.3758/s13421-012-0216-2.

68. Smith MA, Crawford JD (2005). Distributed population mechanism for the 3-D oculomotor reference frame transformation. J Neurophysiol. doi:10.1152/jn.00306.2004

69. Sokal, R.R. and Rohlf, F.J (1987). Introduction to Biostatistics. Freeman, New York.

70. Staub A (2011). The effect of lexical predictability on distributions of eye fixation durations. Psychon Bull Rev. 18(2):371–376. doi:10.3758/s13423-010-0046-9

71. Takahashi M, Shinoda Y (2018). Brain Stem Neural Circuits of Horizontal and Vertical Saccade Systems and their Frame of Reference. Neuroscience. 392:281–328. doi:10.1016/j.neuroscience.2018.08.027

72. Takahashi M, Sugiuchi Y, Na J, Shinoda Y (2022). Brainstem Circuits Triggering Saccades and Fixation. J Neurosci. 42:789–803. doi:10.1523/JNEUROSCI.1731-21.2021

73. Tatler BW, Brockmole JR, Carpenter RH (2017). LATEST: A model of saccadic decisions in space and time. Psychol Rev. 124(3):267–300. doi:10.1037/rev0000054.

74. Treisman AM, Gelade G (1980). A feature-integration theory of attention. Cogn Psychol. 12(1):97–136. doi:10.1016/0010-0285(80)90005-5.

75. Unema PJ, Pannasch S, Joos M, & Velichkovsky BM, (2005). Time course of information processing during scene perception: The relationship between saccade amplitude and fixation duration. Vis Cogn. 12(3), 473–494.

76. Van Gisbergen J and Robinson (1977). Generation of micro and microsaccades by bursts neurons in t h e monkey. In Berthoz (Eds.) Control of Gaze by Brain Stem Neurons, Paris, France R. Baker y A Bertholz. Pp 301–308, Elsevier/North Holland’ Biomedical Press.

77. Van Gisbergen JA, Robinson DA, Gielen S (1981). A quantitative analysis of generation of saccadic eye movements by burst neurons. J Neurophysiol. (3):417–42. doi: 10.1152/jn.1981.45.3.417.

78. Van Gisbergen JA, Van Opstal AJ, Tax AA (1987). Collicular ensemble coding of saccades based on vector summation. Neuroscience. 21:541–555. doi:10.1016/0306-4522(87)90140-0

79. Veale R, Takahashi M (2024). Pathways for Naturalistic Looking Behavior in Primate II. Superior Colliculus Integrates Parallel Top-down and Bottom-up Inputs. Neuroscience.545:86–110. doi:10.1016/j.neuroscience.2024.03.001

80. Takahashi M, Shinoda Y (2018). Brain stem neural circuits of horizontal and vertical saccade systems and their frame of reference. Neuroscience. 392:281–328.

81. Takahashi M, Sugiuchi Y, Na J, Shinoda Y (2022). Brainstem circuits triggering saccades and fixation. J Neurosci. 42:789–803.

82. Takahashi M, Veale R (2023). Pathways for Naturalistic Looking Behavior in Primate I: Behavioral Characteristics and Brainstem Circuits. Neuroscience 532:133–163. doi:10.1016/j.neuroscience.2023.09.009

83. Treisman A, Gelade G. (1980). A feature-integration theory of attention. Cogn Psych. 12, 97–136.

84. Wilming N, Onat S, Ossandón JP, et al. (2017) An extensive dataset of eye movements during viewing of complex images. Sci Data. 4:160126 doi:10.1038/sdata.2016.126

85. White BJ, Berg DJ, Kan JY, Marino RA, Itti L, Munoz DP (2017). Superior colliculus neurons encode a visual saliency map during free viewing of natural dynamic video. Nat Commun. 8:14263 doi:10.1038/ncomms14263

86. Yantis S, Jonides J (1984). Abrupt visual onsets and selective attention: evidence from visual search. J Exp Psychol Hum Percept Perform. 10(5):601–621. doi:10.1037//0096-1523.10.5.601

87. Zee DS, Optican LM, Cook JD, Robinson DA, Engel WK (1976). Slow saccades in spinocerebellar degeneration. Arch Neurol. doi:10.1001/archneur.1976.00500040027004

