## Supplementary material for "Compatibility of a competition model for explaining eye fixation durations during free viewing": figures, statistics, matlab code

### Supplementary figures

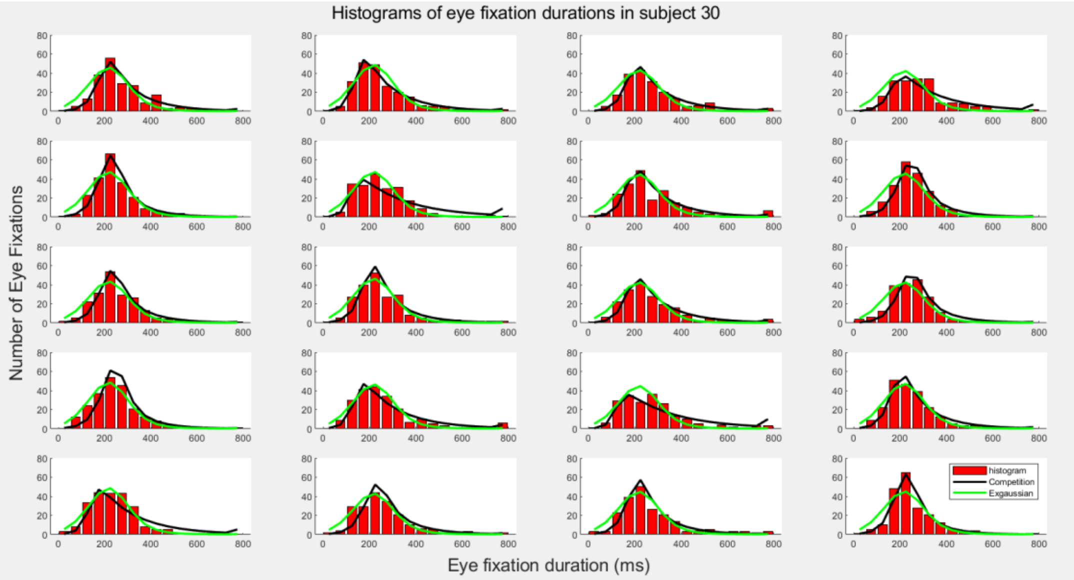


Figure Supplementary 1. *Competition and exgaussian modelling of eye fixation durations histograms. The image shows the fitting of the competition and the exgaussian models of the eye fixation durations frequency histograms for a single subject (subject 30). The fitting is displayed for the four type of images (columns) and for the five block of images presentations (rows).*

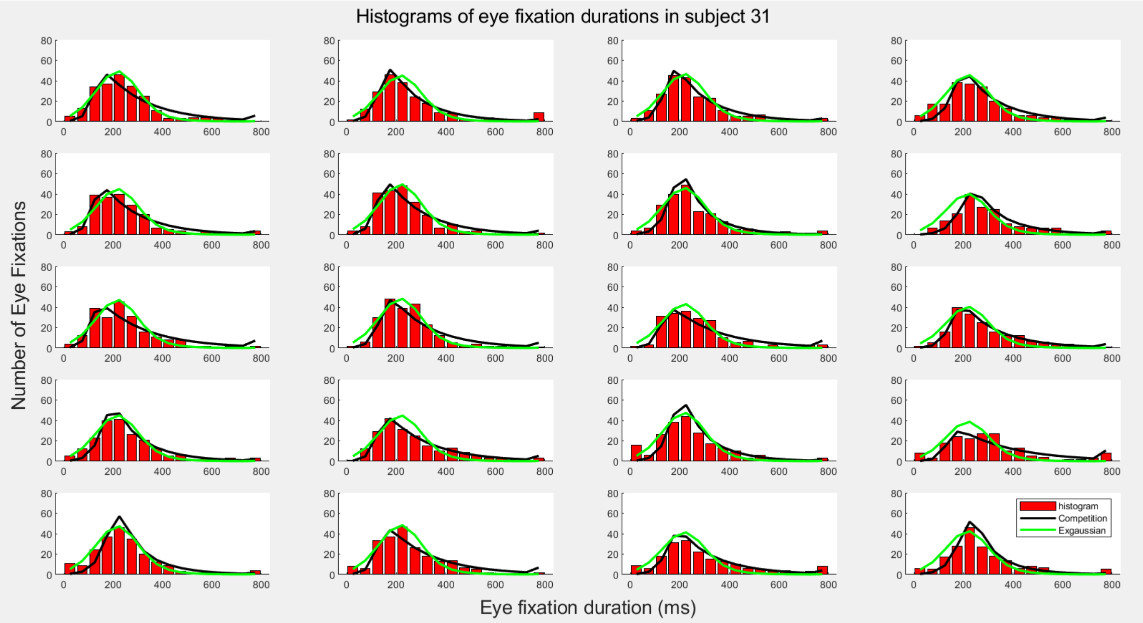


*Figure Supplementary 2. Competition and exgaussian modelling of eye fixation durations histograms. The image shows the fitting of the competition and the exgaussian models of*

*the eye fixation durations frequency histograms for a single subject (subject 31). The fitting is displayed for the four type of images (columns) and for the five block of images presentations (rows).*

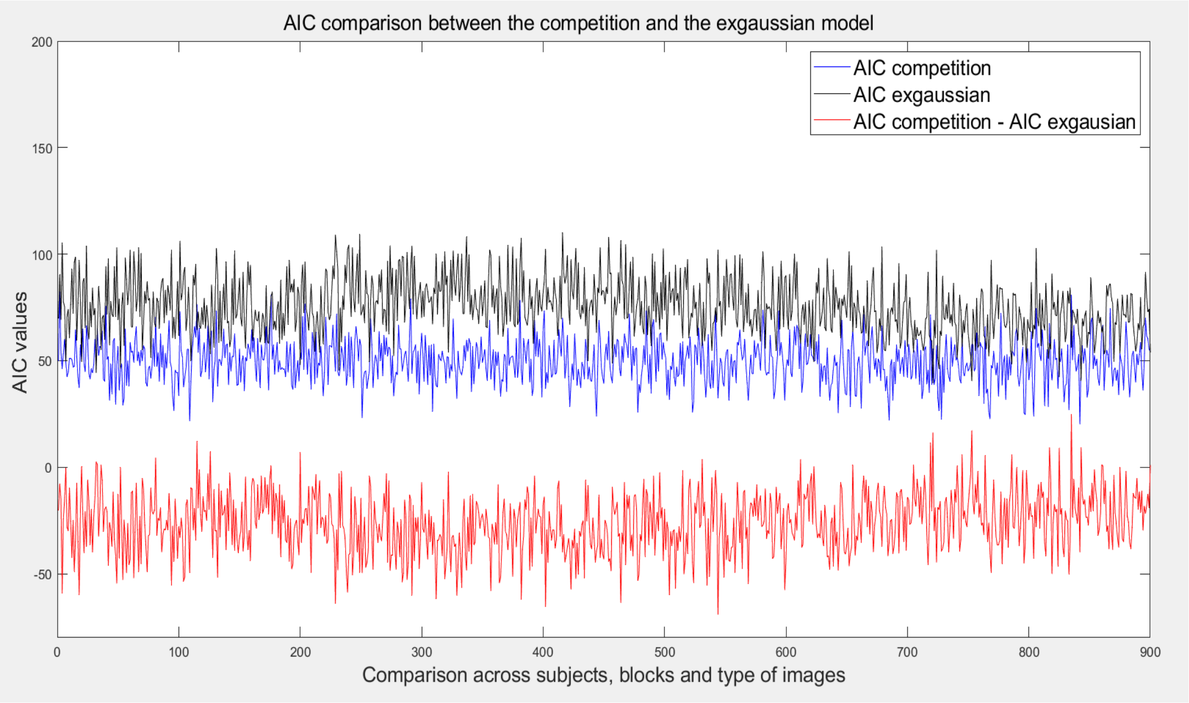


***Supplementary Figure 3, Akaike Information Criteria (AIC).*** *The image shows the AIC values for each of the 900 presented trials (45 subjects, 5 blocks and 4 type of images), for the competition and the exgaussian models. The subtraction of the AIC values of the gausisan model from the AIC values of the competition model are also displayed.*

*Supplementary table 1. Post Hoc Comparisons - block ✻ images*

| | | Mean Difference | SE | df | t | $p_{bonf}$ |
|---|---|---|---|---|---|---|
| B1, nature | B2, nature | -4.327 | 4.121 | 44 | -1.050 | 1.000 |
| | B3, nature | -7.711 | 5.076 | 44 | -1.519 | 1.000 |
| | B4, nature | -10.509 | 5.301 | 44 | -1.983 | 1.000 |
| | B5, nature | -8.638 | 5.343 | 44 | -1.617 | 1.000 |
| | B1, urban | 15.825 | 3.443 | 44 | 4.596 | 0.007 |
| | B2, urban | 10.508 | 3.079 | 44 | 3.413 | 0.264 |
| | B3, urban | 9.661 | 4.902 | 44 | 1.971 | 1.000 |
| | B4, urban | -0.907 | 4.603 | 44 | -0.197 | 1.000 |
| | B5, urban | 0.161 | 5.425 | 44 | 0.030 | 1.000 |
| | B1, fractals | 5.346 | 3.556 | 44 | 1.503 | 1.000 |
| | B2, fractals | -5.013 | 3.957 | 44 | -1.267 | 1.000 |
| | B3, fractals | -9.241 | 4.690 | 44 | -1.970 | 1.000 |
| | B4, fractals | -11.318 | 4.272 | 44 | -2.649 | 1.000 |
| | B5, fractals | -14.739 | 5.720 | 44 | -2.577 | 1.000 |
| | B1, pinknoise | -55.968 | 5.478 | 44 | -10.218 | < .001 |
| | B2, pinknoise | -63.171 | 8.053 | 44 | -7.845 | < .001 |
| | B3, pinknoise | -65.268 | 10.355 | 44 | -6.303 | < .001 |
| | B4, pinknoise | -64.223 | 9.355 | 44 | -6.865 | < .001 |
| | B5, pinknoise | -55.475 | 8.629 | 44 | -6.429 | < .001 |
| B2, nature | B3, nature | -3.384 | 3.435 | 44 | -0.985 | 1.000 |
| | B4, nature | -6.183 | 3.807 | 44 | -1.624 | 1.000 |
| | B5, nature | -4.311 | 3.516 | 44 | -1.226 | 1.000 |
| | B1, urban | 20.152 | 5.500 | 44 | 3.664 | 0.126 |
| | B2, urban | 14.834 | 3.549 | 44 | 4.180 | 0.026 |
| | B3, urban | 13.988 | 4.031 | 44 | 3.470 | 0.223 |
| | B4, urban | 3.420 | 3.622 | 44 | 0.944 | 1.000 |
| | B5, urban | 4.487 | 4.349 | 44 | 1.032 | 1.000 |
| | B1, fractals | 9.673 | 5.457 | 44 | 1.773 | 1.000 |
| | B2, fractals | -0.686 | 4.468 | 44 | -0.154 | 1.000 |
| | B3, fractals | -4.914 | 3.556 | 44 | -1.382 | 1.000 |
| | B4, fractals | -6.991 | 3.901 | 44 | -1.792 | 1.000 |
| | B5, fractals | -10.412 | 5.332 | 44 | -1.953 | 1.000 |
| | B1, pinknoise | -51.641 | 5.128 | 44 | -10.071 | < .001 |
| | B2, pinknoise | -58.844 | 6.802 | 44 | -8.651 | < .001 |
| | B3, pinknoise | -60.942 | 8.051 | 44 | -7.570 | < .001 |
| | B4, pinknoise | -59.896 | 7.220 | 44 | -8.296 | < .001 |
| | B5, pinknoise | -51.148 | 6.198 | 44 | -8.252 | < .001 |
| B3, nature | B4, nature | -2.798 | 3.696 | 44 | -0.757 | 1.000 |
| | B5, nature | -0.927 | 3.762 | 44 | -0.246 | 1.000 |
| | B1, urban | 23.536 | 5.550 | 44 | 4.241 | 0.021 |
| | B2, urban | 18.219 | 3.771 | 44 | 4.832 | 0.003 |
| | B3, urban | 17.373 | 3.135 | 44 | 5.541 | < .001 |
| | B4, urban | 6.804 | 3.726 | 44 | 1.826 | 1.000 |
| | B5, urban | 7.872 | 4.478 | 44 | 1.758 | 1.000 |
| | B1, fractals | 13.057 | 5.987 | 44 | 2.181 | 1.000 |
| | B2, fractals | 2.698 | 4.247 | 44 | 0.635 | 1.000 |

*Supplementary table 1. Post Hoc Comparisons - block ✲ images*

| | | Mean Difference | SE | df | t | $p_{bonf}$ |
|---|---|---|---|---|---|---|
| | B3, fractals | -1.530 | 3.062 | 44 | -0.500 | 1.000 |
| | B4, fractals | -3.607 | 3.760 | 44 | -0.959 | 1.000 |
| | B5, fractals | -7.028 | 4.945 | 44 | -1.421 | 1.000 |
| | B1, pinknoise | -48.257 | 6.012 | 44 | -8.027 | < .001 |
| | B2, pinknoise | -55.460 | 7.098 | 44 | -7.814 | < .001 |
| | B3, pinknoise | -57.557 | 7.584 | 44 | -7.589 | < .001 |
| | B4, pinknoise | -56.512 | 7.339 | 44 | -7.700 | < .001 |
| | B5, pinknoise | -47.764 | 6.127 | 44 | -7.796 | < .001 |
| B4, nature | B5, nature | 1.872 | 2.998 | 44 | 0.624 | 1.000 |
| | B1, urban | 26.335 | 5.982 | 44 | 4.402 | 0.013 |
| | B2, urban | 21.017 | 4.018 | 44 | 5.231 | < .001 |
| | B3, urban | 20.171 | 3.634 | 44 | 5.550 | < .001 |
| | B4, urban | 9.603 | 2.863 | 44 | 3.354 | 0.313 |
| | B5, urban | 10.670 | 3.391 | 44 | 3.147 | 0.563 |
| | B1, fractals | 15.855 | 6.139 | 44 | 2.583 | 1.000 |
| | B2, fractals | 5.496 | 5.095 | 44 | 1.079 | 1.000 |
| | B3, fractals | 1.269 | 3.664 | 44 | 0.346 | 1.000 |
| | B4, fractals | -0.808 | 3.305 | 44 | -0.245 | 1.000 |
| | B5, fractals | -4.230 | 4.144 | 44 | -1.021 | 1.000 |
| | B1, pinknoise | -45.459 | 5.780 | 44 | -7.865 | < .001 |
| | B2, pinknoise | -52.662 | 6.930 | 44 | -7.599 | < .001 |
| | B3, pinknoise | -54.759 | 8.655 | 44 | -6.327 | < .001 |
| | B4, pinknoise | -53.713 | 7.074 | 44 | -7.594 | < .001 |
| | B5, pinknoise | -44.965 | 6.755 | 44 | -6.656 | < .001 |
| B5, nature | B1, urban | 24.463 | 6.224 | 44 | 3.930 | 0.056 |
| | B2, urban | 19.145 | 4.359 | 44 | 4.392 | 0.013 |
| | B3, urban | 18.299 | 4.090 | 44 | 4.474 | 0.010 |
| | B4, urban | 7.731 | 3.306 | 44 | 2.338 | 1.000 |
| | B5, urban | 8.798 | 3.294 | 44 | 2.671 | 1.000 |
| | B1, fractals | 13.983 | 6.439 | 44 | 2.172 | 1.000 |
| | B2, fractals | 3.624 | 5.370 | 44 | 0.675 | 1.000 |
| | B3, fractals | -0.603 | 3.674 | 44 | -0.164 | 1.000 |
| | B4, fractals | -2.680 | 4.261 | 44 | -0.629 | 1.000 |
| | B5, fractals | -6.102 | 4.503 | 44 | -1.355 | 1.000 |
| | B1, pinknoise | -47.331 | 5.545 | 44 | -8.536 | < .001 |
| | B2, pinknoise | -54.533 | 6.567 | 44 | -8.304 | < .001 |
| | B3, pinknoise | -56.631 | 7.602 | 44 | -7.449 | < .001 |
| | B4, pinknoise | -55.585 | 6.940 | 44 | -8.009 | < .001 |
| | B5, pinknoise | -46.837 | 5.224 | 44 | -8.966 | < .001 |
| B1, urban | B2, urban | -5.318 | 3.466 | 44 | -1.534 | 1.000 |
| | B3, urban | -6.164 | 4.432 | 44 | -1.391 | 1.000 |
| | B4, urban | -16.732 | 4.730 | 44 | -3.537 | 0.184 |
| | B5, urban | -15.665 | 5.699 | 44 | -2.749 | 1.000 |
| | B1, fractals | -10.480 | 3.321 | 44 | -3.156 | 0.548 |
| | B2, fractals | -20.839 | 3.933 | 44 | -5.298 | < .001 |
| | B3, fractals | -25.066 | 5.163 | 44 | -4.855 | 0.003 |

*Supplementary table 1. Post Hoc Comparisons - block ✲ images*

| | | Mean Difference | SE | df | t | $p_{bonf}$ |
|---|---|---|---|---|---|---|
| | B4, fractals | -27.143 | 4.850 | 44 | -5.596 | < .001 |
| | B5, fractals | -30.565 | 5.487 | 44 | -5.570 | < .001 |
| | B1, pinknoise | -71.794 | 6.983 | 44 | -10.281 | < .001 |
| | B2, pinknoise | -78.996 | 9.641 | 44 | -8.194 | < .001 |
| | B3, pinknoise | -81.094 | 11.112 | 44 | -7.298 | < .001 |
| | B4, pinknoise | -80.048 | 10.384 | 44 | -7.708 | < .001 |
| | B5, pinknoise | -71.300 | 9.386 | 44 | -7.596 | < .001 |
| B2, urban | B3, urban | -0.846 | 3.127 | 44 | -0.271 | 1.000 |
| | B4, urban | -11.414 | 3.262 | 44 | -3.500 | 0.205 |
| | B5, urban | -10.347 | 4.176 | 44 | -2.478 | 1.000 |
| | B1, fractals | -5.162 | 4.051 | 44 | -1.274 | 1.000 |
| | B2, fractals | -15.521 | 3.343 | 44 | -4.643 | 0.006 |
| | B3, fractals | -19.748 | 3.626 | 44 | -5.447 | < .001 |
| | B4, fractals | -21.825 | 3.010 | 44 | -7.252 | < .001 |
| | B5, fractals | -25.247 | 4.533 | 44 | -5.569 | < .001 |
| | B1, pinknoise | -66.476 | 5.971 | 44 | -11.133 | < .001 |
| | B2, pinknoise | -73.679 | 8.036 | 44 | -9.169 | < .001 |
| | B3, pinknoise | -75.776 | 9.593 | 44 | -7.899 | < .001 |
| | B4, pinknoise | -74.730 | 8.390 | 44 | -8.907 | < .001 |
| | B5, pinknoise | -65.982 | 7.794 | 44 | -8.465 | < .001 |
| B3, urban | B4, urban | -10.568 | 3.074 | 44 | -3.438 | 0.246 |
| | B5, urban | -9.501 | 4.015 | 44 | -2.366 | 1.000 |
| | B1, fractals | -4.316 | 5.093 | 44 | -0.847 | 1.000 |
| | B2, fractals | -14.675 | 3.658 | 44 | -4.012 | 0.044 |
| | B3, fractals | -18.902 | 3.426 | 44 | -5.518 | < .001 |
| | B4, fractals | -20.979 | 3.827 | 44 | -5.482 | < .001 |
| | B5, fractals | -24.401 | 4.063 | 44 | -6.006 | < .001 |
| | B1, pinknoise | -65.630 | 6.341 | 44 | -10.351 | < .001 |
| | B2, pinknoise | -72.832 | 7.922 | 44 | -9.194 | < .001 |
| | B3, pinknoise | -74.930 | 8.883 | 44 | -8.436 | < .001 |
| | B4, pinknoise | -73.884 | 8.229 | 44 | -8.978 | < .001 |
| | B5, pinknoise | -65.136 | 7.165 | 44 | -9.091 | < .001 |
| B4, urban | B5, urban | 1.067 | 3.343 | 44 | 0.319 | 1.000 |
| | B1, fractals | 6.253 | 5.221 | 44 | 1.198 | 1.000 |
| | B2, fractals | -4.107 | 4.285 | 44 | -0.958 | 1.000 |
| | B3, fractals | -8.334 | 3.227 | 44 | -2.582 | 1.000 |
| | B4, fractals | -10.411 | 3.073 | 44 | -3.387 | 0.284 |
| | B5, fractals | -13.832 | 4.244 | 44 | -3.259 | 0.410 |
| | B1, pinknoise | -55.061 | 6.143 | 44 | -8.963 | < .001 |
| | B2, pinknoise | -62.264 | 7.787 | 44 | -7.995 | < .001 |
| | B3, pinknoise | -64.362 | 9.116 | 44 | -7.061 | < .001 |
| | B4, pinknoise | -63.316 | 8.338 | 44 | -7.594 | < .001 |
| | B5, pinknoise | -54.568 | 6.976 | 44 | -7.822 | < .001 |
| B5, urban | B1, fractals | 5.185 | 6.098 | 44 | 0.850 | 1.000 |
| | B2, fractals | -5.174 | 5.741 | 44 | -0.901 | 1.000 |

*Supplementary table 1. Post Hoc Comparisons - block ✲ images*

| | | Mean Difference | SE | df | t | $p_{bonf}$ |
|---|---|---|---|---|---|---|
| | B3, fractals | -9.401 | 4.227 | 44 | -2.224 | 1.000 |
| | B4, fractals | -11.478 | 4.395 | 44 | -2.612 | 1.000 |
| | B5, fractals | -14.900 | 4.012 | 44 | -3.714 | 0.109 |
| | B1, pinknoise | -56.129 | 5.775 | 44 | -9.720 | < .001 |
| | B2, pinknoise | -63.332 | 7.770 | 44 | -8.150 | < .001 |
| | B3, pinknoise | -65.429 | 9.203 | 44 | -7.109 | < .001 |
| | B4, pinknoise | -64.383 | 7.640 | 44 | -8.427 | < .001 |
| | B5, pinknoise | -55.635 | 6.799 | 44 | -8.183 | < .001 |
| B1, fractals | B2, fractals | -10.359 | 3.816 | 44 | -2.715 | 1.000 |
| | B3, fractals | -14.586 | 5.334 | 44 | -2.735 | 1.000 |
| | B4, fractals | -16.664 | 5.072 | 44 | -3.285 | 0.381 |
| | B5, fractals | -20.085 | 5.232 | 44 | -3.839 | 0.075 |
| | B1, pinknoise | -61.314 | 6.687 | 44 | -9.168 | < .001 |
| | B2, pinknoise | -68.517 | 9.343 | 44 | -7.333 | < .001 |
| | B3, pinknoise | -70.614 | 11.203 | 44 | -6.303 | < .001 |
| | B4, pinknoise | -69.568 | 10.391 | 44 | -6.695 | < .001 |
| | B5, pinknoise | -60.820 | 9.704 | 44 | -6.267 | < .001 |
| B2, fractals | B3, fractals | -4.227 | 3.621 | 44 | -1.167 | 1.000 |
| | B4, fractals | -6.305 | 3.633 | 44 | -1.735 | 1.000 |
| | B5, fractals | -9.726 | 4.806 | 44 | -2.024 | 1.000 |
| | B1, pinknoise | -50.955 | 6.486 | 44 | -7.856 | < .001 |
| | B2, pinknoise | -58.158 | 8.537 | 44 | -6.813 | < .001 |
| | B3, pinknoise | -60.255 | 9.362 | 44 | -6.436 | < .001 |
| | B4, pinknoise | -59.209 | 9.491 | 44 | -6.238 | < .001 |
| | B5, pinknoise | -50.461 | 8.281 | 44 | -6.093 | < .001 |
| B3, fractals | B4, fractals | -2.077 | 2.527 | 44 | -0.822 | 1.000 |
| | B5, fractals | -5.499 | 4.199 | 44 | -1.310 | 1.000 |
| | B1, pinknoise | -46.728 | 5.833 | 44 | -8.011 | < .001 |
| | B2, pinknoise | -53.930 | 7.285 | 44 | -7.403 | < .001 |
| | B3, pinknoise | -56.028 | 7.788 | 44 | -7.195 | < .001 |
| | B4, pinknoise | -54.982 | 7.677 | 44 | -7.162 | < .001 |
| | B5, pinknoise | -46.234 | 6.412 | 44 | -7.210 | < .001 |
| B4, fractals | B5, fractals | -3.421 | 4.497 | 44 | -0.761 | 1.000 |
| | B1, pinknoise | -44.650 | 6.074 | 44 | -7.351 | < .001 |
| | B2, pinknoise | -51.853 | 7.869 | 44 | -6.589 | < .001 |
| | B3, pinknoise | -53.951 | 8.753 | 44 | -6.163 | < .001 |
| | B4, pinknoise | -52.905 | 8.000 | 44 | -6.613 | < .001 |
| | B5, pinknoise | -44.157 | 7.504 | 44 | -5.884 | < .001 |
| B5, fractals | B1, pinknoise | -41.229 | 6.139 | 44 | -6.716 | < .001 |
| | B2, pinknoise | -48.432 | 7.691 | 44 | -6.297 | < .001 |
| | B3, pinknoise | -50.529 | 8.898 | 44 | -5.679 | < .001 |
| | B4, pinknoise | -49.483 | 7.844 | 44 | -6.309 | < .001 |
| | B5, pinknoise | -40.735 | 7.149 | 44 | -5.698 | < .001 |
| B1, pinknoise | B2, pinknoise | -7.203 | 6.209 | 44 | -1.160 | 1.000 |

*Supplementary table 1. Post Hoc Comparisons - block ✻ images*

| | | Mean Difference | SE | df | t | $p_{bonf}$ |
|---|---|---|---|---|---|---|
| | B3, pinknoise | -9.300 | 8.372 | 44 | -1.111 | 1.000 |
| | B4, pinknoise | -8.254 | 6.893 | 44 | -1.198 | 1.000 |
| | B5, pinknoise | 0.494 | 6.914 | 44 | 0.071 | 1.000 |
| B2, pinknoise | B3, pinknoise | -2.097 | 7.060 | 44 | -0.297 | 1.000 |
| | B4, pinknoise | -1.052 | 5.502 | 44 | -0.191 | 1.000 |
| | B5, pinknoise | 7.696 | 5.746 | 44 | 1.339 | 1.000 |
| B3, pinknoise | B4, pinknoise | 1.046 | 6.474 | 44 | 0.162 | 1.000 |
| | B5, pinknoise | 9.794 | 5.195 | 44 | 1.885 | 1.000 |
| B4, pinknoise | B5, pinknoise | 8.748 | 5.739 | 44 | 1.524 | 1.000 |

*Note.* P-value adjusted for comparing a family of 190 estimates.

**Script and function to extract the data from the original repository:**

```
% Script to generate the MDTM file for analyzing the data
% First the data must be downloaded from the original database of
% https://datadryad.org/stash/dataset/doi:10.5061/dryad.9pf75
% Then the data of Memory I are extacted using datos =
get_fixmat('etdb_v1.0.hdf5','Memory I');
% the etdb_v1.0.hdf5 and the get_fixmat are obtained from the database
% Now the data must be reorganized to be used by the data analysis script
% enclosed "fitting_competition_exgaussian.m"
% In which the columns includes:
%1. Fixation durations. 2. Subject number.
%3.Order of the figure presentation whithin each subject 4. Block (five
%blocks) 5. Type of figure (four type of blocks)


data = get_fixmat('etdb_v1.0.hdf5','Memory I');
duration_fixation=data.end-data.start;
MDTM=zeros(179473,5);
MDTM(:,1)=duration_fixation;
MDTM(:,2)=data.SUBJECTINDEX;
MDTM(:,3)=data.trial;
MDTM(:,4)=data.iteration;
MDTM(:,5)=data.category;
```

The function required:

```
function fixmat = get_fixmat(database, dataset)
%
%   Read a dataset from the Osnabrueck-Hamburg Eye-Tracking database.
%
%   Returns a struct with fields that encode fixation properties. Each
%   index into a field encodes one fixation.
%
%   Example usage:
%     >> baseline = get_fixmat('etdb_v1.0.hdf5','Baseline')
%
%   Input:
%       database - name of HDF5 file
%       dataset - name of a group within the database file.
%
%   Output:
%       Struct with fields that encode fixation properties.
%
    dataset = sprintf('/%s', dataset);
    info = h5info(database, sprintf('/%s', dataset));
    fixmat = struct();
    for name = {info.Datasets.Name}
        fixmat.(name{1}) = h5read(database, sprintf('%s/%s', dataset,
name{1}));
    end
end
```

**The program for fitting the Competition and the Ex-gaussian model (and the two needed functions):**

```
%%%%%%%%%%%%%%%%%%%%%%%%%%%%%%%%%%%%%%%%%%%%%%%%%%%%%%%%%%%%%%%%%%%%%%%%
% This script (fitting_competition_model) compute the fitting of the
histograms of eye fixation
% durations form the experiment described in the main paper.
%For that it uses the MDTM data file that has reorganized the matrix of eye
fixation
%durations in five fields. 1. Fixation durations. 2. Subject number.
%3.Order of the figure presentation whithin each subject 4. Block (five
%blocks) 5. Type of figure (four type of blocks)
% MDTM It can be requested to the authors or extracted from the
% original data base following the enclosed script "data_extract_MDTM"
% We use two functions: comp_model to compute the C model (created by
% ourselves) and exgfit_1 (modified form exgfit) obtained form other authors
described in the text
%The main resuts are:
%Non_fitting_competition, which give the number of the 900 cases (45
%subjects x 4 types of images x 5 blocks) which don't fit the C model
%Non_fitting_ex_gaussianidem for for the
%exgaussian model

load('MDTM.mat')
```

```
clearvars -except MDTM


close all
hist_values=0;
intervalos=[25:50:1425];
threshold = 1500;
global_cuentas=0,
cuentas=0;
s = [100,100,100];
for sujeto=1:1:45
sujeto
    n=1;
    for bloque=1:1:5
        for tipo_imagen=1:1:4
            clear values
            if tipo_imagen==1
                numeracion_imag=7;
                values = MDTM(MDTM(:,2) == sujeto & MDTM (:,4)== bloque &
MDTM (:,5)==7);
                % Remove elements in the vector that exceed the threshold
                values1=values;
                values(values > threshold) = [];
                mean_values=mean(values);
                mean_global(sujeto,bloque,tipo_imagen)=mean_values;
                std_values=std(values);
                std_global(sujeto,bloque,tipo_imagen)=std_values;
                nn=size(values);
                nn=nn(1,1);
                nn_global(sujeto,bloque,tipo_imagen)=nn;
                cuentas=size(values1);
                global_cuentas=global_cuentas + cuentas;
                hist_values=hist(values,intervalos);
                %plot (intervalos, hist_values);
            end

            if tipo_imagen==2
                numeracion_imag=8;
                values =MDTM( MDTM(:,2) == sujeto & MDTM (:,4)== bloque &
MDTM (:,5)==8);
                % Remove elements in the vector that exceed the threshold
                values1=values;
                values(values > threshold) = [];
                mean_values=mean(values);
                mean_global(sujeto,bloque,tipo_imagen)=mean_values;
                std_values=std(values);
                std_global(sujeto,bloque,tipo_imagen)=std_values;
                nn=size(values);
                nn=nn(1,1);
                nn_global(sujeto,bloque,tipo_imagen)=nn;
                cuentas=size(values1);
                global_cuentas=global_cuentas + cuentas;
                hist_values=hist(values,intervalos);

            end
            if tipo_imagen==3
                numeracion_imag=10;
                values = MDTM( MDTM(:,2) == sujeto & MDTM (:,4)== bloque &
MDTM (:,5)==10);
```

```
                % Remove elements in the vector that exceed the threshold
                values1=values;
                values(values > threshold) = [];
                mean_values=mean(values);
                mean_global(sujeto,bloque,tipo_imagen)=mean_values;
                std_values=std(values);
                std_global(sujeto,bloque,tipo_imagen)=std_values;
                nn=size(values);
                nn=nn(1,1);
                nn_global(sujeto,bloque,tipo_imagen)=nn;
                cuentas=size(values1);
                global_cuentas=global_cuentas + cuentas;
                hist_values=hist(values,intervalos);

            end
            if tipo_imagen==4
                numeracion_imag=11;
                values = MDTM(MDTM(:,2) == sujeto & MDTM (:,4)== bloque &
MDTM (:,5)==11);
                % Remove elements in the vector that exceed the threshold
                values1=values;
                values(values > threshold) = [];
                mean_values=mean(values);
                mean_global(sujeto,bloque,tipo_imagen)=mean_values;
                std_values=std(values);
                std_global(sujeto,bloque,tipo_imagen)=std_values;
                nn=size(values);
                nn=nn(1,1);
                nn_global(sujeto,bloque,tipo_imagen)=nn;
                cuentas=size(values1);
                global_cuentas=global_cuentas + cuentas;
                hist_values=hist(values,intervalos);

            end

            [f4, correlacion, tj, A_out, p_out,
h,p_kolm,ks2stat,f2_max,hist_values_mod,f4_mod] = comp_model(hist_values);
            correlacion_global(sujeto,bloque,tipo_imagen)=correlacion;
            hist_values_global(sujeto,bloque,:,tipo_imagen)=hist_values;

hist_values_global_mod(sujeto,bloque,:,tipo_imagen)=hist_values_mod;
            f2_max_global(sujeto,bloque,:,tipo_imagen)=f2_max;
            f4_global(sujeto, bloque,:,tipo_imagen)=f4;
            f4_global_mod(sujeto, bloque,:,tipo_imagen)=f4_mod;
            A_out_global(sujeto,bloque,tipo_imagen)=A_out;
            p_out_global(sujeto,bloque,tipo_imagen)=p_out;
            h_global(sujeto,bloque,tipo_imagen)=h;
            p_kolm_global(sujeto,bloque,tipo_imagen)=p_kolm;
            ks2stat_global(sujeto,bloque,tipo_imagen)=ks2stat;

            [mu,sigma,tau] = exgfit_1(values,s);
             mu_global(sujeto,bloque,tipo_imagen)=mu;
            sigma_global(sujeto,bloque,tipo_imagen)=sigma;
            tau_global(sujeto,bloque,tipo_imagen)=tau;
            n=n+1

        end
    end
end
```

```
mu_mean = mean(mean(mean(mu_global, 1), 2), 3); % Mean across all dimensions
sigma_mean = mean(mean(mean(sigma_global, 1), 2), 3); % Mean across all
dimensions
tau_mean = mean(mean(mean(tau_global, 1), 2), 3); % Mean across all
dimensions
%to obtain the numebr of cases which does not fit the model

Non_fitting_competition=sum(h_global,'all');


%Now for fitting the exgaussian
%optimization of exgaussian parameters  by iteration

            mu_i=fix(mu_mean)-100;
            mu_f=fix(mu_mean)+100;
            sigma_i=fix(sigma_mean)-50;
            sigma_f=fix(sigma_mean)+50;
            tau_i=fix(tau_mean)-50;
            tau_f=fix(tau_mean)+50;


for sujeto=1:1:45;

    sujeto
    for bloque=1:1:5
        for tipo_imagen=1:1:4

            p_max=0;
            for mu=mu_i:5:mu_f
                for sigma=sigma_i:5:sigma_f
                    for tau=tau_i:5:tau_f
                        for i=1:1:29

                            % Define the Ex-Gaussian PDF
                            %mu=mu_global(sujeto,bloque,tipo_imagen);
                            %sigma=sigma_global(sujeto,bloque,tipo_imagen);
                            %tau=tau_global(sujeto,bloque,tipo_imagen);% Ex-
Gaussian Probability Density Function
                            % x     - Points where PDF is evaluated
                            % mu    - Mean of the Gaussian component
                            % sigma - Standard deviation of the Gaussian
component
                            % tau   - Scale parameter of the exponential
component

                            % Compute the Ex-Gaussian PDF
                            term1 = 1 ./ tau;
                            term2 = exp((sigma^2 ./ (2 * tau^2)) -
(((intervalos(i)) - mu) ./ tau));
                            term2=squeeze(term2);
                            term3 = 0.5 * erfc((1 / sqrt(2)) * ((sigma / tau)
- (((intervalos(i)) - mu) ./ sigma)));
                            term3=squeeze(term3);


                            % Final Ex-Gaussian PDF
                            ex_f2(i) = term1 .* term2 .* term3;
                            normalization=1/(sum(ex_f2));
                            ex_f2=ex_f2*normalization;
```

```
                            %compute the frequency histogram of fixations
durations
                            n=nn_global(sujeto,bloque,tipo_imagen);
                            ex_f4=ex_f2*n;

                        end
                        %fitting exgaussian to histogram of frequency
durations


hist_values_mod=hist_values_global_mod(sujeto,bloque,:,tipo_imagen);
                        hist_values_mod=squeeze(hist_values_mod);
                        hist_values_mod=hist_values_mod';
                        ex_f4_mod(1,1:16)=zeros;
                        ex_f4_mod(1,1:15)=ex_f4(1,1:15);
                        ex_f4_mod(1,16)=sum(ex_f4(1,16:29));
                        [h,p_kolm,ks2stat] =
kstest2(hist_values_mod,ex_f4_mod);

ex_f4_global_mod(sujeto,bloque,:,tipo_imagen)=ex_f4_mod;
                        ex_h_global(sujeto,bloque,tipo_imagen)=h;
                        ex_p_kolm_global(sujeto,bloque,tipo_imagen)=p_kolm;
                        ex_ks2stat_global(sujeto,bloque,tipo_imagen)=ks2stat;
                        mu_global(sujeto,bloque,tipo_imagen)=mu;
                        sigma_global(sujeto,bloque,tipo_imagen)=sigma;
                        tau_global(sujeto,bloque,tipo_imagen)=tau;
                        %keeping the statistics
                        if p_kolm>p_max

ex_f4_global_mod(sujeto,bloque,:,tipo_imagen)=ex_f4_mod;
                        ex_h_global(sujeto,bloque,tipo_imagen)=h;
                        ex_p_kolm_global(sujeto,bloque,tipo_imagen)=p_kolm;
                        ex_ks2stat_global(sujeto,bloque,tipo_imagen)=ks2stat;
                        mu_global(sujeto,bloque,tipo_imagen)=mu;
                        sigma_global(sujeto,bloque,tipo_imagen)=sigma;
                        tau_global(sujeto,bloque,tipo_imagen)=tau;
                        p_max=p_kolm;
                        end

                    end
                end
            end
        end
    end
end

Non_fitting_ex_gaussian=sum(ex_h_global,'all');
```

**The two required functions:**

```
%computing the best fitting using the competition model
```

```
function [f4, correlacion, tj, A_out, p_out,
h,p_kolm,ks2stat,f2_max,hist_values_mod,f4_mod ] =
comp_model(hist_values)
% This function returns the sum and product of two numbers
%pruebas
%data=[1:1:30];
%p=0.2;
%A=0.3;
%%%%%%%%%%%
clear A_out p_out A p f f1 f2 f2_max f3 f4 correlacion_1 sum_data
size_dat size_data prueba hist_values_mod f4_mod
sum_data=sum(hist_values);
size_dat=size(hist_values);
size_data=size_dat(1,2);
hist_values=hist_values';
contador=1;


%compute the best p and A parameters


%close all
correlacion=0;
tj=1:1:size_data;
for p=0.01:0.01:0.99
    for  A=0.1:0.1:4.9
        for t=1:1:size_data;


            f(t,1)=((1-p)^(t-1))*p;
            t1=(-1)*t;
            f1(t,1)=1/(1+(exp((A*t1)+exp(2))));
            f2=f1*(p);
            %f3=1-f3;
            f3(t,1)=((1-f2(t))^(t-1))*f2(t);


        end
        prob_sum=sum(f3);
        norm1=1/prob_sum;
        f3=f3*norm1;
        correlacion_1=corr (hist_values,f3);
        %correlacion_1
        if correlacion_1 > correlacion
            correlacion_out=correlacion_1;
            correlacion=correlacion_out;
            A_out=A;
            p_out=p;
            maximo=contador;
        end
        contador=contador+1;
        prueba(contador,:)=f3;
```

```
        prueba_1(contador,:)=f2;
    end
end




f4=prueba(maximo,:)*sum_data;
f2_max=prueba_1(maximo,:);
%[h,p_kolm,ks2stat] = kstest2(hist_values,f4);
% same but collapsing values higher than 1000
hist_values=hist_values';
hist_values_mod(1,16)=0;
hist_values_mod(1,1:15)=hist_values(1,1:15);
hist_values_mod(1,16)=sum(hist_values(1,16:29));
f4_mod(1,1:16)=zeros;
f4_mod(1,1:15)=f4(1,1:15);
f4_mod(1,16)=sum(f4(1,16:29));
[h,p_kolm,ks2stat] = kstest2(hist_values_mod,f4_mod);
end
```

```
% exgfit  Fit ExGaussian distribution to data
%
% [MU,SIGMA,TAU] = exgfit(X,S) fits the ExGaussian distribution to data in
% vector X using maximum likelihood and returns the fitted parameters MU,
% SIGMA, and TAU. The ExGaussian distribution is formed by the sum of
% independent normal and exponential observations. MU and SIGMA denotes
% the mean and standard deviation of the normal component and TAU denotes
% the mean of the exponential component. S is a three-element vector of
% starting values for MU, SIGMA, and TAU when fitting the distribution to
% data. SIGMA must be positive and TAU at least zero.
%
%   Example
%         n = 200;
%         mu = 500; sigma = 200; tau = 400;
%         x = randn(1,n)*sigma+mu-log(1-rand(1,n))*tau;
%         s = [100,100,100];
%         [mu,sigma,tau] = exgfit(x,s);
%         % Below plots the results without Statistics Toolbox
%         ncdf = @(x) 0.5*(1+erf(x/sqrt(2)));
%         epdf = @(x,mu,sigma,tau) (1/tau).*exp((mu/tau)+(sigma^2/(2*tau^2))-
...
%             (x/tau)).*ncdf( (x-mu-(sigma^2/tau))./sigma);
%         histogram(x,'Normalization','pdf','Facecolor',[.6,.6,.6]);
%         hold on
%         xx = linspace(min(x),max(x),1e3);
%         plot(xx,epdf(xx,mu,sigma,tau),':k','linew',2);
%         set(gca,'Color',[.98,.98,.98],'FontSize',15)
%         grid on
%         xlabel('X','FontSize',25)
%         ylabel('pdf','FontSize',25)
```

```
function [mu,sigma,tau] = exgfit_1(x,s)
ncdf = @(x) 0.5*(1+erf(x/sqrt(2)));

epdf = @(x,mu,sigma,tau) (1/tau).*exp((mu/tau)+(sigma^2/(2*tau^2))-...
    (x/tau)).*ncdf( (x-mu-(sigma^2/tau))./sigma);
L = @(p) sum(-log(epdf(x,p(1),p(2),p(3))));
%original fit = fmincon(L,s,[],[],[],[],[-Inf,0+eps,0],[]);
%quasi original [mu,sigma,tau] = deal(pfit(1),pfit(2),pfit(3));
% Define the objective function (negative log-likelihood)
L = @(p) sum(-log(epdf(x, p(1), p(2), p(3))));

% Set the tolerance for stopping criteria
options = optimoptions('fmincon', ...
    'Display', 'iter', ...       % Display iteration information
    'MaxIter', 1000, ...         % Maximum number of iterations
    'MaxFunEvals', 10000, ...    % Maximum function evaluations
    'OptimalityTolerance', 1e-8, ... % Tolerance for first-order optimality
(gradient)
    'StepTolerance', 1e-8, ...   % Tolerance for step size change between
iterations
    'FunctionTolerance', 1e-8, ... % Tolerance for change in function value
between iterations
    'Algorithm', 'interior-point'); % Algorithm choice, can also try 'trust-
region-reflective'

%options = optimoptions('fmincon', 'MaxIter', 1000, 'MaxFunEvals', 10000,
'Display', 'iter');
pfit = fmincon(L, s, [], [], [], [], [0+eps, 0+eps, 0+eps], [400, 400, 400],
[], options);
[mu,sigma,tau] = deal(pfit(1),pfit(2),pfit(3));
end
```

**Script for simulating the competition model and generate figure 1:**

```
% This script simulate the competition dynamics of the model described in
% the manuscript


clear all
close all


% Parameters
n = 500; % Number of points in each time series
rng(11); % Set seed for reproducibility


% Create two overlapping distributions for the time series
ts1 = 4 + 2 * randn(n, 1); % Time series 1 (μ=4, σ=2)
ts2 = 6 + 2 * randn(n, 1); % Time series 2 (μ=6, σ=2)
```

```
% Ensure they are uncorrelated by orthogonalizing
ts2 = ts2 - (ts1'*ts2)/(ts1'*ts1) * ts1;


% Normalize to maintain original distribution properties
ts1 = (ts1 - mean(ts1))/std(ts1) * 2 + 4;
ts2 = (ts2 - mean(ts2))/std(ts2) * 2 + 6;


% Create figure with horizontal layout
figure('Position', [100, 100, 1200, 500], 'Color', 'White');


%% First subplot: Time series with upward crossings
subplot(3, 2, 1);
plot(1:n, ts1, 'b', 'LineWidth', 1.5);
hold on;
plot(1:n, ts2, 'r', 'LineWidth', 1.5);


% Detect only upward crossings (TS1 crosses above TS2)
diff_series = ts1 - ts2;
sign_changes = diff(sign(diff_series));
upward_crossings = find(sign_changes == 2);  % Only positive crossings
upward_crossings =upward_crossings+1;
% Plot upward crossings with green triangles
scatter(upward_crossings, ts1(upward_crossings), 50, 'g', 'filled', '^');
title('A) Time Series of networks activity');
xlabel('Time');
ylabel('Activity');
legend('Saccade network', 'Eye fixation network', 'Upward Crossings',
'Location', 'best');
grid on;


%% Second subplot: Time between upward crossings
subplot(3, 2, 2);


if length(upward_crossings) > 1
    time_between = diff(upward_crossings);
    histogram(time_between, 'BinMethod', 'integers', 'FaceColor', [0.5
0.5 0.5]);
    hold on;
    %xline(mean(time_between), 'r--', 'LineWidth', 2, ...
         % 'Label', ['Mean: ' num2str(mean(time_between), 2)]);
    title('B) Time Between Upward Crossings');
    xlabel('Time Steps Between Crossings');
    ylabel('Frequency');
    grid on;
else
```

```
    text(0.5, 0.5, 'No upward crossings detected', 'HorizontalAlignment',
'center');
    title('No Upward Crossings Found');
end





%comutation of geometric



%figure('color','white');



for p=0.25:0.1:0.7


    nf=100;
    j=1
    for t=1:0.1:10
        f(j)=((1-p)^(t-1))*p;
        j=j+1
    end
    tt=[0:5:450];
    a=sum(f);
    % normalization of the histogram to make to integral of f1=1
    f1=f/a;
    b=sum(f1)
    f1=f1*nf
    % Plot the first subplot
    subplot(3, 2, 3);
    plot(tt,f1);
    hold on
    title('C) Histogram of eye fixations durations  \newline by geometric
pdf (eq. 1), changing ps');
    xlabel('Eye fixation durations (ms)');
    ylabel('N of fixations');
end
hold off



% Dependency of the saccadic-latency network probability to win
competition from the time
```

```
%elapsed from previous saccade to make the sigmoid asyntotically approach
to the value of saccade winning
%the competition (ps)
p=0.3;
A=5;
j=1;
for t=1:0.1:10
    t1=(-1)*t;
    f2(j)=(1/(1+(exp((A*t1)+exp(2)))));
    j=j+1
end


f2=f2*(p);
f3=1-f2;




subplot(3, 2, 4);
plot(tt,f2);
hold on
plot(tt,f3);
title('D) Probability of fixation (1-ps`(t)) and saccadic  \newline
network (ps`(t))to win competition (eq.3)');
xlabel('time from previous saccade');
ylabel('ps` and (1-ps`)' );
legend('saccadic network (ps`)','fixation network (1-ps`)',
'location','best')



% Complete competition model changing A parameter keeping ps
fila=1;


p=0.1
for A=2:1:6
    j=1
    for t=1:0.2:10
        t1=(-1)*t;
        f2(fila,j)=(1/(1+(exp((A*t1)+exp(2)))));
        j=j+1
    end
    fila=fila+1;
end
f2=f2*(p);


for fila=1:1:5
    j=1
```

```
    for t=1:1:46
        f4(fila,t)=((1-f2(fila,t))^(t-1))*f2(fila,t);

    end
end


a=sum(f4,2);
b=repmat(a,1,46);


% normalization of the histogram to make to integral of f1=1
f5=f4./b;
nf=100;
f6=f5*nf;
% computing the histogram of fixation times
ttt=[0:10:450];
subplot(3, 2, 5);
plot(ttt,f6);
title('E) Histogram of of eye fixations durations  \newline changing
parameter A (eq. 2))');
    xlabel('Eye fixation durations (ms)');
    ylabel('N of fixations');








% Complete competition model changing A parameter keeping ps
fila=1;
A=3;


for p=0.02:0.06:0.35
    j=1
    for t=1:0.2:10
        t1=(-1)*t;
        f7(fila,j)=(1/(1+(exp((A*t1)+exp(2)))))*p;
        ttt(j)=j;
        j=j+1
    end
    fila=fila+1;
end


for fila=1:1:5
    j=1
    for t=1:1:46
        f8(fila,t)=((1-f7(fila,t))^(t-1))*f7(fila,t);
        ttt(j)=j;
```

```
    end
end


a=sum(f8,2);
b=repmat(a,1,46);


% normalization of the histogram to make to integral of f1=1
f9=f8./b;


nf=100;
f10=f9*nf;
% computing the histogram of fixation times
ttt=[0:10:450];
subplot(3, 2, 6);
plot(ttt,f10);
title('F) Histogram of of eye fixations durations  \newline changing
parameter ps (eq. 1 and 3))');
    xlabel('Eye fixation durations (ms)');
    ylabel('N of fixations');
```

**Script for generating the data histograms and the fitting of the competition and exgaussian model:**

```
close all
%display subject by subject the  histogram and the fitted models
figure('color','white');
hold on
close all
%tttj=(tj*50)-25;
ttj=tj(1:16);
ttj=(ttj*50)-25

selected_subject=18

n=1;


for bloque=1:1:5
    for tipo_imagen=1:1:4
        subplot (5,4,n);


        subplot (5,4,n);
        zz=hist_values_global_mod(selected_subject,bloque,:,tipo_imagen);
        zz=squeeze(zz);
        zzz=f4_global_mod(selected_subject,bloque,:,tipo_imagen);
```

```
        zzz=squeeze(zzz);
        zzzz=ex_f4_global_mod(selected_subject,bloque,:,tipo_imagen);
        zzzz=squeeze(zzzz);
        %plot(ttj,zz,'r',ttj,zzz,'k',ttj,zzzz,'g','LineWidth', 1);
        hold on; % Hold the current plot
bar(ttj, zz, 'r'); % Plot zz as bars (histogram-style)
plot(ttj, zzz, 'k', 'LineWidth', 2); % Plot zzz as a line
plot(ttj, zzzz, 'g', 'LineWidth', 2); % Plot zzzz as a line
hold off; % Release the hold
        ylim([0 80]);
        if bloque==5 & tipo_imagen==4;
        legend('histogram', 'Competition ','Exgaussian' ); % Adding legend
        end
        n=n+1
        hold on
    end
end
han = axes(gcf, 'Visible', 'off');
han.XLabel.Visible = 'on';
han.YLabel.Visible = 'on';

xlabel(han, 'Eye fixation duration (ms)', 'FontSize', 16);
ylabel(han, 'Number of Eye Fixations ','FontSize', 16);
%sgtitle({'Histograms of Fixation Durations in subject 1'},'FontSize', 16); %
Common title

sgtitle({sprintf('Histograms of eye fixation durations in subject %d',
selected_subject),}, 'FontSize', 16);
hold off
```
